# *In Vivo* Pathway Optimization in Yeast via *LoxPsym*-Mediated Shuffling of Upstream Activating Sequences

**DOI:** 10.1101/2024.12.21.629127

**Authors:** Ciaran Ruehmkorff, Ali Tafazoli Yazdi, Lena Hochrein

## Abstract

The budding yeast *Saccharomyces cerevisiae* plays an integral role in the bioeconomy as a powerful host for industrial bio-manufacturing, driving the production of diverse bio-based products. Achieving optimal product yields requires precise fine-tuning of the expression levels of multiple pathway genes, which often relies on cloning-intensive methods. Here, we present PULSE, an *in vivo* promoter engineering tool based on a streamlined workflow combining FACS-based screening of a randomized DNA library to identify active promoter elements, and their subsequent assembly into synthetic hybrid promoters where each element is flanked by *loxPsym* sites. Multiple promoter cassettes can be genome-integrated to generate “ready-to-use” platform strains, allowing users to easily place their genes of interest under the control of PULSE promoters. By activating Cre-mediated recombination, *loxPsym*-flanked promoter elements can be recombined, effectively bringing the target genes under control of a vast set of promoters spanning a wide range of expression levels in one simple step. Applying PULSE on two heterologous pathways, an eightfold increase in β-carotene production and improved growth on high xylose concentrations by *S. cerevisiae* was achieved. These results demonstrate the power and efficiency of PULSE as a versatile platform for metabolic engineering, enabling rapid, cloning-free optimization of biosynthetic pathways *in vivo*.

## Introduction

The budding yeast *Saccharomyces cerevisiae* is widely used in industrial applications due to its fast growth, robust fermentation capabilities, well-studied genetics, GRAS status and ease of genetic manipulation (1). With its ability to be easily scaled up and its tolerance for industrial conditions, it is considered as an ideal host organism to produce bulk and fine chemicals such as biofuels, nutraceuticals and natural products (2, 3). The microbial synthesis of such compounds often requires the introduction of heterologous metabolic pathways, which typically consist of multiple enzymatic steps. To increase product yields, it is crucial to avoid disturbances to metabolic homeostasis and to optimize the metabolic flux through the non-native pathway. This requires fine-tuning the expression levels of both heterologous and native genes, as precise control of key enzymes significantly enhances cell growth and product formation (4, 5). Promoters are key elements for regulating transcriptional levels, making them a powerful tool to fine-tune enzyme expression and optimize the efficiency and stability of metabolic pathways. Given that overexpression of all genes in a pathway is not a guaranteed success for enhancing product formation and predicting the optimal promoter strengths remains a challenge, the optimal combination of promoter expression must be experimentally determined for each biosynthetic pathway (6). For this, *in vitro* combinatorial assembly strategies based on Type IIS restriction enzymes or homology regions are commonly employed to generate large libraries in which each gene in a pathway is regulated by different promoter variants (7–10). However, these approaches have two major drawbacks. First, repeatedly using the same promoter can lead to unintended recombination events, potentially decreasing strain stability and productivity. To mitigate this issue, these methods rely on a large set of well-characterized promoters that cover a broad range of expression levels. Second, the generation of combinatorial libraries is often restricted by the transformation efficiency of the host organism and typically requires cost- and labor-intensive cloning procedures. Given these limitations, forming a promoter library directly *in vivo* offers a more practical alternative.

Cuperus et al. (2015) demonstrated the first *in vivo* method to fine-tune the expression of three lycopene biosynthesis genes in *S. cerevisiae* (11). To modulate promoter strength, they employed six mutant tetracycline operator (*tetO*) sites with varying affinities for the synthetic transcription factor TetR-VP16. These *tetO* variants were initially introduced to *S. cerevisiae* on a plasmid and subsequently released from the plasmid using a homing endonuclease. This allowed the *tetO* sequences to integrate randomly into the yeast genome through homologous recombination, positioning themselves upstream of minimal promoters regulating the target genes. This results in a synthetic promoter library providing a range of expression levels regulating the pathway genes. However, the study reports that only approximately 0.7% of the engineered yeast population successfully produced detectable levels of target product lycopene, indicating successful genome-integration and expression of pathway genes. The low efficiency significantly increases the number of strains that need to be screened, requiring high-throughput methods and additional resources to identify the best producers, which limits scalability and practicality without further optimization.

The Cre-*loxP* recombination system provides another powerful tool for the generation of *in vivo* libraries which has been demonstrated in the Sc2.0 project (12). Cre-mediated recombination of *loxPsym* sites can lead to deletion, inversion, translocation and duplication of DNA fragments with approximately equal probability (13). In their toolkit GEMbLeR, Cautereels et al. (2024) developed a hybrid promoter and terminator each consisting of six native regulatory elements flanked by orthogonal *loxPsym* sites. The recombination sites enable the independent rearrangement of the order and orientation of the native promoter and terminator elements. This promoter-terminator cassette was then used to regulate each of the six genes required for the synthesis of astaxanthin. Strong promoters resulting from recombination mostly depend on the very strong *TDH3* promoter element, potentially limiting the diversity of expression levels (14). Li et al. (2025) developed a similar version of this tool based on native promoter elements with the focus on establishing a recombination system with an increased inversion rate (15). While these tools succeed at generating *in vivo* promoter libraries they heavily depend on specific elements such as the WT-*tetO* (11) or *TDH3* promoter element (14, 15) to reach a strong expression level and therefore show drawbacks in their modularity. Further, repeated usage of the same promoter elements could impact genomic stability.

Our promoter shuffling tool PULSE **(P**romoter engineering by shuffling **U**pstream activating sequences via ***L****oxPsym* **S**upported **E**volution) overcomes these limitations by providing a streamlined three-step workflow to generate and identify diverse promoter cassettes tailored for *loxPsym*-mediated recombination. Starting with a simplified FACS-based screening of a randomized DNA library from a single yeast transformation, we identified numerous active upstream activating sequence (UAS) elements. These elements were assembled into hybrid promoter cassettes that achieved expression levels exceeding those of the strong native *TDH3* promoter. By integrating multiple promoter cassettes into the yeast genome, platform strains that provide a robust foundation for pathway optimization can be created, enabling simultaneous, dynamic, and cloning-free fine-tuning of the expression levels of multiple genes *in vivo* based on the Cre-*loxP* system. In PULSE, each pathway gene is regulated by a distinct synthetic hybrid promoter, composed of a core promoter element and unique combinations of UAS elements flanked by *loxPsym* sites. Cre activity alters the order, orientation and number of UAS elements and thus the strength of the promoter. In this way, starting from an isogenic yeast clone, we generated a culture of diverse yeast strains spanning a wide range of expression levels post-recombination, independent of cloning and transformation efficiency.

Applying the PULSE system for the optimization of heterologous pathway expression, we achieved a β-carotene production of 70 mg/L, representing a nearly eightfold increase over a reference strain already equipped with strong, unrecombined PULSE promoters. Furthermore, by leveraging PULSE to the xylose utilization pathway, we improved the growth of *S. cerevisiae* under high xylose concentrations, evidenced by a reduced lag-phase and increased final biomass, demonstrating the versatility of PULSE across distinct metabolic contexts. Importantly, these optimizations were achieved by testing less than 100 candidates, demonstrating the efficiency and potency of the PULSE tool in rapidly identifying improved strains.

## Material and Methods

### Strains and media

*Escherichia coli* strains DH5α, NEB5α and DB3.1 (New England Biolabs, Frankfurt am Main, Germany) were used for cloning and grown at 37°C and 230 rpm in Luria-Bertani (LB) medium with antibiotic selection, *i.e.* carbenicillin (100 µg/mL), chloramphenicol (34 µg/mL), or kanamycin (50 µg/mL). All generated yeast strains (listed in Supplementary Table 1A) are derivates of the *S. cerevisiae* strain BY4742 (16) and were cultured at 30°C and 220 rpm in Yeast extract Peptone Dextrose Adenine (YPDA)-rich medium or in Complete Supplement Mixture (CSM) medium.

### Plasmid construction using the MoClo-YTK toolkit

For plasmid assembly the *S. cerevisiae* MoClo toolkit (Addgene: Kit #1000000061) (17) was extensively used, which provides a modular and standardized assembly system, enabling the rapid construction of plasmids through Golden Gate cloning. DNA parts such as promoters, coding sequences (CDS), terminators as well as markers and origins of replication are hierarchically assembled into entry plasmids, which define their final position in the target vector. Primers used throughout this study are listed in Supplementary Table 1B. As part of this work, we generated several new part plasmids and added them to the MoClo toolkit. To enable the assembly of hybrid promoters, the promoter part (part 2) was divided into two subparts: 2a, containing UAS elements, and 2b, containing the core promoter. The nucleotide sequence GTAA was chosen as the new Golden Gate Assembly overhang between the 2a and 2b parts (18). Additionally, to facilitate the genomic integration of single markerless gene cassettes, five homology regions from the EasyClone-MarkerFree toolkit (X-3, XI-2, XII-2, XII-5, XI-3) (19) were adapted and added as MoClo parts 1 and 5, replacing the connective linkers. An alternative MoClo entry plasmid (pYTK_UP_22p) was also developed, which uses a *CcdB* gene instead of *GFP* to allow counter-selection. The *CcdB* gene can be easily replaced with a new MoClo part in a Golden Gate assembly with Esp3I. All part plasmids created in this study are listed in Supplementary Table 2.

The completed MoClo parts library consisting of both, existing and newly generated parts, was used to assemble the following plasmids: The UAS entry plasmid pYTK_UP_001c, which holds a *loxPsym* flanked *CcdB* spacer upstream of the minimal Core 1 promoter (20) regulating *yEGFP* expression, served as the backbone for the insertion of the UAS library. Its expression cassette is flanked by homology regions for genome-integration into the yeast XII-2 locus (19). To allow genomic integration of gene cassettes into the five distinct genomic sites, five entry plasmids, pYTK_UP_003c, 009c and 013c-015c, were generated, which hold the required homology regions and a *CcdB* spacer, that allows the later integration of a gene cassette as parts 2−4 (Supplementary Table 3). These plasmids were utilized to assemble the native promoter constructs regulating *yEGFP* and *mScarlet* expression in the different genomic loci via BsaI Golden Gate assemblies. Details about the assembled plasmids can be retrieved from Supplementary Table 4.

### Generation of the randomized UAS library

The UAS library (pAT06) was generated by first converting the oligonucleotide CR_147 containing 48 bases of randomized nucleotides flanked by fixed sequences of 24 and 36 nucleotides, into double-stranded DNA through a single DNA amplification cycle. The primer CR_018 anneals to 18 bp of the fixed region of CR_147. A DNA amplification involving three steps starting with an initial denaturation at 98°C for 15 s, followed by 10 s of annealing at 55°C, and finishing with a final step of elongation for 100 min at 68°C was performed. Five ng of the resulting double-stranded amplicons were then digested with restriction enzymes KpnI and NheI and ligated into 50 ng of the digested pYTK_UP_001c backbone in a 16 µL reaction, to exchange the *CcdB* gene for members of the UAS library (Supplementary Figure 1A and 1B upper part). This resulted in a plasmid library holding 48 bp of randomized sequence, flanked by restriction enzyme recognition sites as well as *loxPsym* sites in the following order: SacI, SalI, *loxPsym*, NheI, Esp3I (upstream) and BsaI, KpnI, *loxPsym* and HindIII (downstream). The negative UAS control (pAT11) was assembled following the same principle as pAT06, except for using CR_182 instead of CR_147 as the oligonucleotide.

### Assembly of multi-UAS plasmids

The multi-UAS plasmids were assembled by specific restriction and ligation reactions assigning each UAS element to a predefined position (Supplementary Figure 1B; Supplementary Table 5). To assemble a 3xUAS promoter, three single UAS plasmids were digested in a predefined pattern. Overhangs of Type IIS restriction enzymes were designed in a way that digestion with Esp3I generates a fitting sticky end overhang to the HindIII overhang and digestion with BsaI generates a fitting overhang to SalI. The UAS elements directly upstream of the core promoter were marked as position 1 (P1). The P1 UAS plasmids were digested with SacI and Esp3I to serve as backbone into which two smaller fragments derived from other UAS plasmids were ligated as P2 (digested with SalI and HindIII) and P3 (digested with SacI and BsaI). The 3xUAS promoter could now be further expanded to harbor five UAS elements. To this end, the 3xUAS plasmid needed to be digested with SalI and HindIII to serve as the middle fragment in the 5xUAS promoter. A single, distinct UAS element is inserted upstream and another distinct UAS element with the whole plasmid backbone is placed downstream of the pre-assembled triple UAS cassette, allowing the promoter cassette to be expanded by two UAS elements at a time.

### Cloning multi-UAS promoters with the K528 Kozak sequence

The K528 Kozak sequence (GCAATA) (21) was added downstream of the core promoter of the multi-UAS plasmids via HiFi assembly (Supplementary Table 6). Here the whole multi-UAS promoter plasmids were amplified with primers CR_635 and L819 giving homology to the region flanking the Kozak sequence (22 bp upstream and 20 bp downstream), which was supplied in the form of the annealed oligonucleotides CR_636/CR_637.

### Cloning of the Δ*loxPsym* promoter cassettes

To exchange the *loxPsym* sites in the 11xUAS element (pCR_343), a gene fragment was synthesized (Twist Bioscience, South San Francisco, USA) in which the twelve *loxPsym* sites were replaced by twelve unique 34 bp sequences. These sequences were selected based on a predicted minimal number of transcription factor binding motifs, as determined using the YeTFaSCo webtool (22). The gene fragment was PCR-amplified using different primer pairs to generate UAS cassettes of varying lengths (Supplementary Table 7). The resulting PCR fragments were inserted into the backbone of pCR_191, containing the Core 1 promoter regulating *yEGFP* expression, via restriction and ligation cloning.

### Cloning of 5xUAS cassettes with single UAS at different positions

To assemble the 5xUAS cassettes with single UAS elements at the three distinct positions, a series of pre plasmids had to be assembled. First, to enable later assemblies, a BbsI site had to be removed from the nUAS-K528 containing plasmid (pCR_191) by assembling two PCRs (template pCR_191, primers CR_755/CR_660 and CR_754/CR_666) in a HiFi reaction to form pCR_276. Next, a *loxPsym*-flanked *CcdB* gene was generated by amplifying *CcdB* with primers CR_756 and CR_757 from plasmid pYTK_UP_001c and digest it with PaqCI and BbsI before ligating it into the BsaI and Esp3I digested plasmid pCR_276. Using these plasmids as building blocks, a 3xnUAS promoter was assembled (pCR_278), as well as a nUAS-*CcdB*-nUAS promoter following the multi-UAS assembly strategy (Supplementary Table 5). These 3xUAS plasmids were furthermore expanded to form 5xUAS promoter cassettes using specified restriction enzyme digestions followed by ligation, which is described in more detail in Supplementary Table 8. In this way, three new UAS-entry plasmids (pCR_280– 282) were constructed, each containing a *CcdB* gene at positions 1, 3 or 5, with nUAS sequences occupying the remaining positions. These backbones were subsequently digested with PaqCI and BbsI to remove the *CcdB* gene, and the UAS elements were inserted as annealed oligonucleotides (Supplementary Table 9).

### Assembly of final PULSE plasmids

Final PULSE plasmids were assembled by first generating two levels of pre plasmids. In the first level, five UAS-CDS entry plasmids (pCR_420–424) were assembled using the MoClo toolkit. Each plasmid was assigned for a different genome-integration site (X-3, XI-2, XII-2, XII-5, XI-3)(19) and furthermore harbored gene cassettes with a *CcdB* gene as part 2a, the core 1 promoter (20) with the K528 Kozak (21) sequence as part 2b, a further *CcdB* gene at position 3 and the *SPG5*-terminator as part 4 (Supplementary Table 10). The *CcdB* genes in part 2a of pCR_420–424 were exchanged for the PULSE promoters 5xUAS_A–E in a digestion and ligation reaction described in detail in Supplementary Table 11 to form the PULSE-CDS entry plasmids. The remaining *CcdB* gene in part 3 of the PULSE-CDS entry plasmids can be excised by PaqCI digestion and exchanged for a desired CDS in a HiFi reaction. The CDSs were either PCR amplified (for *yEGFP*, *mScarlet* and the β-carotene genes) or directly used as a gene fragment (for the xylose oxidoreductase pathway genes), both with 20–25 bp homology overhangs to the PULSE-CDS entry plasmid backbones. A detailed assembly plan of all assembled PULSE plasmids is depicted in Supplementary Table 12. The homology regions were designed in a way that there is a seamless transition in between the K528 Kozak sequence and the CDS, with no additional nucleotides in between.

### Cloning of the 5xUAS_C-pTDH3-core promoter cassette

To assemble the 5xUAS_C-pTDH3-core promoter, the pAT07 plasmid harboring the *TDH3* promoter was amplified as a backbone with primers CR_1031 and CR_1032. Subsequently this PCR product and plasmid pCR_231, harboring the 5xUAS-C cassette, were digested with SacI and HindIII and ligated to form pCR_444, harboring the 5xUAS-C-pTDH3-core promoter.

### Plasmid isolation and cell transformation

Commercially available chemically competent *E. coli* strains NEB5α (New England Biolabs, Frankfurt am Main, Germany) and DB3.1 (Thermo Fisher Scientific, Waltham, MA, USA), or homemade competent DH5α cells were transformed following the High Efficiency Transformation Protocol of New England Biolabs. Plasmids were extracted from *E. coli* using the NucleoSpin Plasmid EasyPure kit (Machery-Nagel, Düren, Germany) and verified by sequencing (LGC Genomics, Berlin, Germany).

To generate the UAS library, NEB5α cells were transformed with 16 µL of the ligation reaction and plated onto 14 LB plates with carbenicillin. This resulted in a total of 6,200 colonies which were washed from the plates and combined into a single vial, followed by plasmid isolation.

Yeast transformations were performed using the LiAc/SS carrier DNA/PEG method by Gietz and Schiestl (23) using 500 ng of plasmid DNA. Genomic integration of gene cassettes into specified loci was done using the homology regions X-3, XI-2, XII-2, XII-5 and XI-3 provided by the CRISPR/CAS9-based EasyClone-MarkerFree toolkit with their respective helper plasmids (19). Gene cassettes flanked by the homology regions were amplified using the PrimeSTAR^®^ MAX polymerase (Takara Bio, Saint-Germain-en-Laye, France) with the primers CR_315/CR_316 for X-3; CR_564/CR_565 for XI-2; CR_564/CR_565 for XI-2; CR_305/CR_314 for XII-2; CR_311/CR_312 for XII-5 and CR_309/CR_310 for XI-3. Integration into a single locus was done by co-transformation of 700 ng of linearized DNA fragments with a helper plasmid expressing one sgRNA or in three loci at ones with a plasmid expressing three sgRNAs for X-3, XI-2 and XII-2.

### Fluorescence-activated cell sorting

Yeast liquid precultures were inoculated from a single colony taken from a YPDA/CSM-URA agar plate in 500 µL of CSM/CSM-URA in 48-well microtiter plates (Greiner, Frickenhausen, Germany). The precultures were incubated at 30°C with 220 rpm shaking for 30 h. Precultures were then used to inoculate main cultures to an OD_600_ of 0.05 in 500 µL of CSM/CSM-URA in the 48-well microtiter plates. The main cultures were grown at 30°C and 220 rpm shaking for approximately 13 h for measurements in the logarithmic growth phase and for 30 h for measurements in the stationary growth phase.

Yeast samples were treated with 500 µg/mL cycloheximide (Merck, Darmstadt, Germany) to stop protein production before FACS analysis. The cells were filtered through a 40 µm cell strainer (pluriSelect, Leipzig, Germany) and the fluorescence output of single cells was measured with the BioRad S3e cell sorter (BioRad Laboratories, Munich, Germany) using 100 mW lasers with wavelengths of 488 nm for yEGFP and 561 nm for mScarlet. The yeast population was gated in a forward scatter to side scatter dot plot and 50,000 cells were counted per sample at a speed of 2,000 events/s. The fluorescence height was analyzed in a histogram and the geometrical mean was calculated for each measurement using the Flowlogic software (Miltenyi Biotec B.V. & Co. KG, Bergisch Gladbach, Germany). Each sample was normalized either to the *PFY1* promoter or the *TDH3* promoter integrated into the XII-2 locus or to mScarlet/mCherry fluorescence regulated by the *ADH1*/*CYC1* promoter for episomal samples.

To sort the randomized UAS library, the transformed yeast colonies were washed from the transformation plates with CSM-URA and pooled to form strain Y1433. Four mL of CSM-URA in a 25-mL shake flask were inoculated with 10 µL of the Y1433 strain and grown overnight. Fluorescence was analyzed with the 488 nm laser. A sorting gate was designed including the top 2% of the *yEGFP* expressing cells in the library. A total of 200,000 cells were sorted from this gate at a flow rate of 2,000 events/s. The sorted cells were plated on CSM-URA agar plates and incubated for 3 days at 30°C.

To sort the Cre-recombined PULSE promoter libraries (Y2006, Y2008, Y2010, Y2012 and Y2014) (details in section “Light-induced Cre-recombinase experiments”), the yeast libraries were inoculated to an OD_600_ of 0.05 in 10 mL CSM in a shake flask after light treatment and incubated at 30°C with 220 rpm shaking for 30 h, before they were reinoculated as a main culture under the same conditions again. The cells grew until they reached the exponential growth-phase and were then kept for 10 h in the logarithmic growth phase without exceeding the OD_600_ of 1. The cells were washed and resuspended in ice-cold PBS and kept on ice until sorting. Fluorescence was analyzed with the 488 nm laser. Each sample was divided into ten uniform bins ranging from the lowest to the highest fluorescence output of the sample. In each bin 10,000 cells were sorted at a speed of 2,000 events/s and subsequently grown in 500 µL of CSM for 30 h in a 48-well microtiter plate (Greiner, Frickenhausen, Germany). The recombined yeast promoters in each bin were PCR amplified using PrimeSTAR^®^ GXL DNA Polymerase (Takara Bio) with the appropriate primer pair (Supplementary Table 13) and the promoter size was analyzed on a 2% agarose gel. Furthermore, the binned yeast cultures were reinoculated and analyzed by FACS.

### Pretesting UAS elements in a microplate reader

A total of 744 single colonies from the CSM-URA plates, obtained after FACS, were inoculated into 250 µL of liquid CSM-URA in 96-well microtiter plates (Sarsted, Nuembrecht, Germany) as precultures and incubated overnight at 30°C with 220 rpm shaking. The main cultures were inoculated in 250 µL liquid CSM-URA medium in 96-well microtiter plates to an OD_600_ of 0.1 and grown at 30°C with 220 rpm shaking for 30 h to reach the stationary phase. The yEGFP and mScarlet signals were measured using an Infinite^®^ 200 PRO microplate reader (Tecan, Maennedorf, Swiss). The gain was adjusted by the plate reader based on the average signal of all samples on the plate. To neglect the effect of multiple plasmid copies in a single cell, the yEGFP signal was normalized using the mScarlet signal, since a mScarlet expression cassette under the control of the *ADH1* promoter was present on each plasmid. The normalized yEGFP fluorescence was used to determine active UAS elements. Samples showing more than two times the signal strength of the negative control (nUAS) were considered as active UAS elements.

### Growth analysis of strains containing multi-UAS promoter cassettes

Yeast liquid precultures were inoculated from single colonies taken from a YPDA/CSM-URA agar plate into 500 µL of CSM in 48-well microtiter plates (Greiner, Frickenhausen, Germany). The precultures were incubated at 30°C with 220 rpm shaking for 30 h. Precultures were then used to inoculate main cultures to an OD_600_ of 0.05 in 500 µL of CSM in the 48-well microtiter plates (Greiner, Frickenhausen, Germany). In the initial growth experiment, which included the BY4742 wildtype strain as well as strains harboring PULSE promoters, native promoters or 1-13xUAS cassettes, main cultures were grown at 30°C with shaking at 220 rpm shaking for 30 h. Optical density (OD_600_) was measured at multiple time points using an Infinite^®^ 200 PRO microplate reader (Tecan, Maennedorf, Swiss).

Growth rates (µ) were calculated according to the formula µ=(ln(OD_2_) – ln(OD_1_)) / (t_2_ - t_1_) where OD_1_ and OD_2_ are the optical densities at time points t_1_ and t_2_, respectively. The lag-phase duration was determined using the tangent method, in which the end of the lag-phase is defined by the intersection of the baseline (initial biomass) with the tangent of the maximal growth rate µ_max_ (24, 25). Doubling times during exponential growth phase were calculated using the equation doubling time = ln(2) / µ_max_.

In the second growth experiment including the BY4742 wildtype as well as strains harboring multi-nUAS promoters or multi-UASΔloxPsym promoters, main cultures were cultivated in 200 µl of CSM in black 96-well microtiter plates with transparent bottom (Porvair Science Ltd, Norfolk, UK) with 4 mm orbital shaking in the Infinite^®^ 200 PRO microplate reader for 30 h at 30°C and OD_600_ measurements every 15 min.

### Light- induced Cre-recombinase experiments

Yeast cultures were inoculated to an OD_600_ of 0.05 in 500 µL CSM-LEU containing 25 µM PCB (phycocyanobilin, SiChem GMBH, Bremen, Germany) in 24-well plates with a transparent bottom (product no. 303008, Porvair Science Ltd, Norfolk, UK) sealed with sterile aluminum sealing foil. The plates were transferred into a custom-made light plate apparatus device (26). The device was equipped with a 660 nm LED (product no. L2-0-R5TH50-1, LEDsupply, Randolph, VT, USA) and a 740 nm LED (product no. MTE1074N1-R, Marktech Optoelectronics Inc., Latham, NY, USA). The cultures were treated with a 30-s far-red light pulse and subsequently grown for 8 h at 30°C and 220 rpm in darkness. Subsequently, the cultures were irradiated with a single 30-s red light pulse, followed by 10-s red light pulses every 5 min over the indicated period, while uninduced cells were cultured in the dark. The induction period ended with a 1 min far-red light pulse (740 nm).

### Oxford Nanopore sequencing of PULSE promoters

Genomic DNA of the recombined yeast cultures (strains Y2006, Y2008, Y2010, Y2012 and Y2014) was extracted using the YeaStar^©^ Genomic DNA kit (Zymo Research, Freiburg, Germany). The recombined promoters derived from the 5xUAS_A–E promoters were amplified using the KAPA HiFi HotStart ReadyMix PCR kit (Roche, Basel, Schweiz) with appropriate primer pairs (Supplementary Table 13). The PCR products were purified using Agencourt^©^ AMPure^©^ XP beads (Beckmann Coulter, Brea, USA) and analyzed on a 2% agarose gel. Further, the amplicons were prepared for Nanopore sequencing according to the Ligation Sequencing DNA V14 (SQK-LSK114) protocol provided by Oxford Nanopore Technologies (Oxford, UK). The library was loaded onto two Flongle flowcells (FLO-FLG114) (Oxford Nanopore Technologies, Oxford, UK). The sequencing runs lasted 24 h and generated 197,000 and 461,000 reads, with an N50 of 609 and 637 base pairs, respectively. Base calling of the Nanopore sequencing run was performed by the “High-accuracy model v4.3.0, 400 bp”. The sequencing results were pooled and used for further analysis. The pooled sequencing data was processed by a custom program made in python utilizing the Biopython library (27) to quantify the distribution of the resulting strains after recombination (Supplementary Tables 14–18). First, each read was aligned to the list of EasyClone genomic locus sequences (X-3, XI-2, XII-2, XII-5 and XI-3) using the pairwise alignment tool provided within the Biopython library. Next, each sequence was aligned to a list of all the isolated UAS elements. The resulting analysis reported the number, position, and orientation (second number after the UAS identity, 1 = normal orientation, 2 = inversion) of the UAS elements for each read along with the corresponding construct in a CSV file. Subsequently, this data was used to identify unique UAS arrangements and determine their frequency for each promoter construct. This information revealed the total diversity of promoters and their relative abundances after recombination.

### DNA quantification via agarose gel electrophoresis

PCR amplicons on a gel picture were quantified using the Quantity tool of the Image Lab™ desktop software Version 6.0.1 (BioRad Laboratories, Munich, Germany). The molar amount of each fragment was then calculated using the “Mass -> Moles” section in the NEBioCalculator^®^ (https://nebiocalculator.neb.com/#!/dsdnaamt). The amount of mols of each PCR amplicon was then divided by the total amount of mols to receive the percentage of each amplicon of the total DNA.

### Growth experiment over 100 generations of the β-PULSE-0 strain

The β-PULSE-0 strain was cultivated in 20 mL of CSM medium using the Chi.Bio turbidostat platform (28) in dither mode, starting at an initial OD_600_ of 0.1. In this setup, cells were repeatedly grown to an OD_600_ of ∼0.6, at which point they were diluted back to an OD_600_ of 0.3. Each of these growth-dilution cycles was defined as one generation. The reactor was operated for more than 200 h until more than 100 dilution steps had occurred. The culture was then re-streaked on a CSM agar plate, and genomic DNA was extracted from both the β-PULSE-0 preculture and the culture after the passage of 100 generations. The four PULSE promoters were amplified using the KAPA HiFi HotStart ReadyMix PCR kit (Roche, Basel, Schweiz) with the appropriate primer pairs (Supplementary Table 13). The PCR products were purified using Agencourt*©* AMPure*©* XP beads (Beckmann Coulter, Brea, USA) and analyzed on a 2% agarose gel. Further, the amplicons were prepared for Nanopore sequencing according to the Ligation Sequencing DNA V14 (SQK-LSK114) protocol provided by Oxford Nanopore Technologies (Oxford, UK). The library was loaded onto a Flongle flowcell (FLO-FLG114) (Oxford Nanopore Technologies, Oxford, UK). The sequencing runs lasted 24 h and generated 360,000 reads, with an N50 read length of 958 base pairs. Base calling of the Nanopore sequencing run was performed by the Fast model (v4.3.0) with a target speed of 400 bp/s. The data processing and analysis was performed as described in the “Oxford Nanopore sequencing of PULSE promoters” section.

### β-Carotene quantification

For the pre-tests in small volume, yeast liquid precultures were inoculated from a single colony taken from a CSM agar plate in 500 µL of CSM in a 48-well microtiter plate (Greiner, Frickenhausen, Germany). The precultures were incubated at 30°C with 220 rpm shaking for 30 h before they were inoculated to an OD_600_ of 0.05 in 500 µL of CSM in 48-well microtiter plates as a main culture and grown at 30°C and 220 rpm for 72 h. Afterwards, the cells were transferred to a 2-mL safelock Eppendorf tube, spun down at 13,000 *g* for 5 min and washed with ddH_2_O. 200 µL of glass beads with 0.5 mm diameter, and 1 mL acetone were added to the tube. The samples were treated for 5 min at 30 Hz in the MM400 mixer mill (Retsch, Haan, Germany), subsequently incubated for 20 min at 30°C and then centrifuged for 5 min at 16700 *g* at 4°C. The clear supernatant was filtered through a 0.22 µm PTFE filter into a glass vial.

For the final tests in shake flasks, yeast liquid precultures of β-PULSE-0 and β-PULSE-1DY were inoculated from a single colony taken from a CSM agar plate in 500 µL of CSM in a 48-well microtiter plate (Greiner, Frickenhausen, Germany). Precultures were incubated at 30°C with 220 rpm shaking for 30 h, before being used to inoculate main cultures to an initial OD_600_ of 0.05 in 200 mL CSM medium in 2 L baffled flasks. Main cultures were grown at 30°C and 220 rpm for 242 h. At multiple time points, the OD_600_ was measured, and 5 mL of culture were transferred to 15-mL Falcon tubes for β-carotene extraction. The cultures were spun down at 3,000 *g* for 5 min, resuspended in 1 mL of ddH_2_O, transferred to 2-mL safelock Eppendorf tubes and spun down again at 13,000 *g* for 5 min. 500 µL of glass beads with 0.5 mm diameter and 1 mL acetone were added to the tube. The samples were treated for 5 min at 30 Hz in an MM400 mixer mill (Retsch, Haan, Germany), subsequently incubated for 20 min at 30°C, and then centrifuged for 5 min at 16700 *g* at 4°C. 700 µL the clear supernatant was filtered through a 0.22-µm PTFE filter into a glass vial and 700 µL of fresh acetone was added to the extraction tube. The extraction process was restarted beginning with the MM400 mixer mill and repeated for four rounds in total.

A Nexera HPLC (Shimadzu, Berlin, Germany) was used, equipped with a Symmetry C18 column 100 Å, 5 µm, 4.6 mm X 250 mm (Waters, Milford, USA) for reversed phase chromatography. For β-carotene detection, an isocratic program with acetone:methanol:isopropanol (85:10:5 v/v/v) was employed as a mobile phase with a flow of 1 mL/min for 10 min at 25°C (29). The β-carotene peak was analyzed at 454 nm and had a retention time of 4.8 min. The lycopene peak was detected at 474 nm with a retention time of 4 min. Calibration curves were generated using purified standards (β-carotene at concentrations of 1, 10, 50 and 200 mg/L; and lycopene at 0.1, 1, and 10 mg/L) to quantify compound concentration in the extracts.

### RT-qPCR

The RNA of three biological replicates of the strains β-PULSE-0, β-PULSE-1DY, X-PULSE-0 and X-PULSE-G11, harvested in the exponential and the stationary growth phase, was extracted using the YeaStar^TM^ RNA kit (Zymo Research, Freiburg, Germany). A DNase treatment was performed using the TURBO-DNA-free^TM^ kit (Thermo Fisher Scientific, Waltham, MA, USA). Subsequently, cDNA was amplified with the RevertAid H Minus First Strand cDNA Synthesis Kit (Thermo Fisher Scientific, Waltham, MA, USA) and utilized as a template for the RT-qPCR. The primer pairs and conditions for each gene are listed in Supplementary Table 19. The TAF10 gene was used as the reference gene to determine the ΔCT value and the PULSE-0 strains were used as the reference strains for determining the ΔΔCT values.

### Whole genome sequencing of β-PULSE-1DY

Genomic DNA extraction and whole genome sequencing of β-PULSE-1DY was performed by Plasmidsaurus (Cologne, Germany) using Oxford Nanopore Technology with custom analysis and annotation. An amplification-free, long-read sequencing library was generated using the v14 library prep chemistry, including minimal fragmentation of the gDNA in a sequence independent-manner. This library was sequenced using a primer-free protocol on R10.4.1 flow cells, generating 143,141 reads with an estimated coverage of 90x and an N50 of 806,925 base pairs.

### Xylose growth experiments

After light induction in CSM-LEU with 20 g/L glucose, 250 µL of the cells were inoculated in 10 mL of CSM with either 10 g/L xylose or 50 g/L xylose in a shake flask and grown at 30°C with 220 rpm shaking for 30 h and subsequently recultured under the same conditions. 3,000 cells of each culture were plated on a vented QTray agar plate (Molecular Devices, San José, USA) with CSM (50 g/L xylose). Forty-seven colonies originating from the 10 g/L xylose preculture and 47 colonies originating from the 50 g/L xylose preculture were randomly picked into a 96-deepwell plate containing 1 mL of CSM (50 g/L xylose) using the QPix 420 colony picker (Molecular Devices, San José, USA). The cultures were incubated for five days at 30°C and 800 rpm in an Infors Multitron plate shaker (Axon Labortechnik, Kaiserslautern, Germany) and reinoculated to an OD_600_ of 0.05 and grown under the same conditions as the preculture. After 0, 13, 20 and 36 h samples were taken for OD_600_ measurements in the Infinite^®^ 200 PRO plate reader. Fifteen strains that grew fast and reached a high cell density were selected for the next test round. These strains and the unrecombined X-PULSE strain as a control were inoculated in three technical replicates in a 48 well plate to an OD_600_ of 0.05 in 500 µL CSM (50 g/L xylose). The strains were grown in the Infinite^®^ 200 PRO plate reader at 30°C with orbital shaking (4 mm) for 48 h with an OD_600_ measurement every 15 min. In a final screening the four best strains and the control (X-PULSE-0) were inoculated to an OD_600_ of 0.05 in 500 µL of CSM with different xylose concentrations (5, 10, 30 and 50 g/L xylose) in 48-well plates in three independent biological replicates with three technical replicates. The plates were incubated at 30°C with 220 rpm shaking for 82 h and the OD_600_ was measured at 13 timepoints throughout the exponential and stationary growth phases using the Infinite^®^ 200 PRO plate reader.

### Xylose, xylitol and ethanol quantification

Three biological replicates of X-PULSE-0 and X-PULSE-G11 were cultivated at an initial OD_600_ of 0.05 in 50 mL of CSM containing 50 g/L xylose in 500 mL baffled flasks. At a series of designated time points, 1 mL samples were collected, and the OD_600_ of the cultures was determined. Then, each sample was centrifuged for 3 min at 20,000 x g and 540 µL of the supernatant was transferred to a new Eppendorf tube and mixed with 50 % (w/v) 5-sulfosalycilsäure (SSA), vortexed, and incubated for 5 min at RT before a second centrifugation step at 20,000 x g for 5 min. Xylose, xylitol and ethanol amounts in the supernatant were then quantified by Dr. Mislav Oreb (Goethe University Frankfurt) using HPLC, as described in Regmi et al. (2024) (30). Growth rates were calculated with the formula µ=(ln(OD_2_) – ln(OD_1_)) / (t_2_ - t_1_). The lag-phase duration was determined using the tangent method, where the end of the lag-phase is marked by the intersection of the initial biomass with a line tangent of µ_max_ (24, 25).

## Results and Discussion

This study aimed to develop a versatile promoter engineering toolkit for fine-tuning the expression of multiple genes *in vivo*, enhancing bioproduction or growth on alternative carbon sources. Our PULSE toolkit is based on hybrid promoter cassettes that allow dynamic adaptation of gene expression levels through Cre-mediated recombination of UAS elements. Analogous to native yeast promoters, our hybrid promoters are divided into a core promoter and UAS elements located upstream of the core promoter (31, 32). While the core promoter is responsible for the recruitment and assembly of the preinitiation complex and the initiation of transcription, UAS elements serve as binding sites for specific transcription factors and affect promoter activity (33). Our promoter cassettes consist of at least five synthetic UAS elements positioned upstream of a previously characterized minimal Core 1 promoter (20), yielding high strength promoter cassettes. Each UAS element is flanked by symmetrical *loxPsym* sites, enabling Cre mediated recombination to generate deletions, inversions, insertions, or duplications of individual UAS elements (Figure 1A) (12). In this way, the PULSE tool enables a single promoter cassette to be converted into a diverse library of promoter variants *in vivo* that differ in the size, composition and orientation of their UAS elements, resulting in a broad spectrum of promoter strengths. Finally, these cassettes are integrated into the yeast genome to generate platform strains for optimization of any pathway of interest (Figure 1B). To optimize flux through a pathway, users simply place the coding sequences (CDS) of target genes under the control of PULSE promoters. Activation of the Cre recombinase simultaneously diversifies multiple promoter cassettes starting from a single isogenic strain, generating a yeast library with a range of promoter strengths regulating pathway genes. This enables fine-tuning of gene expression to achieve the optimal ratio of proteins, enhancing pathway flux and strain productivity. Importantly, this process is independent of the host organism’s cloning or transformation efficiency. Ideal candidates with improved traits, such as higher product yield or growth advantages, can be identified and isolated from the library through high-throughput screening.

**Figure 1.**
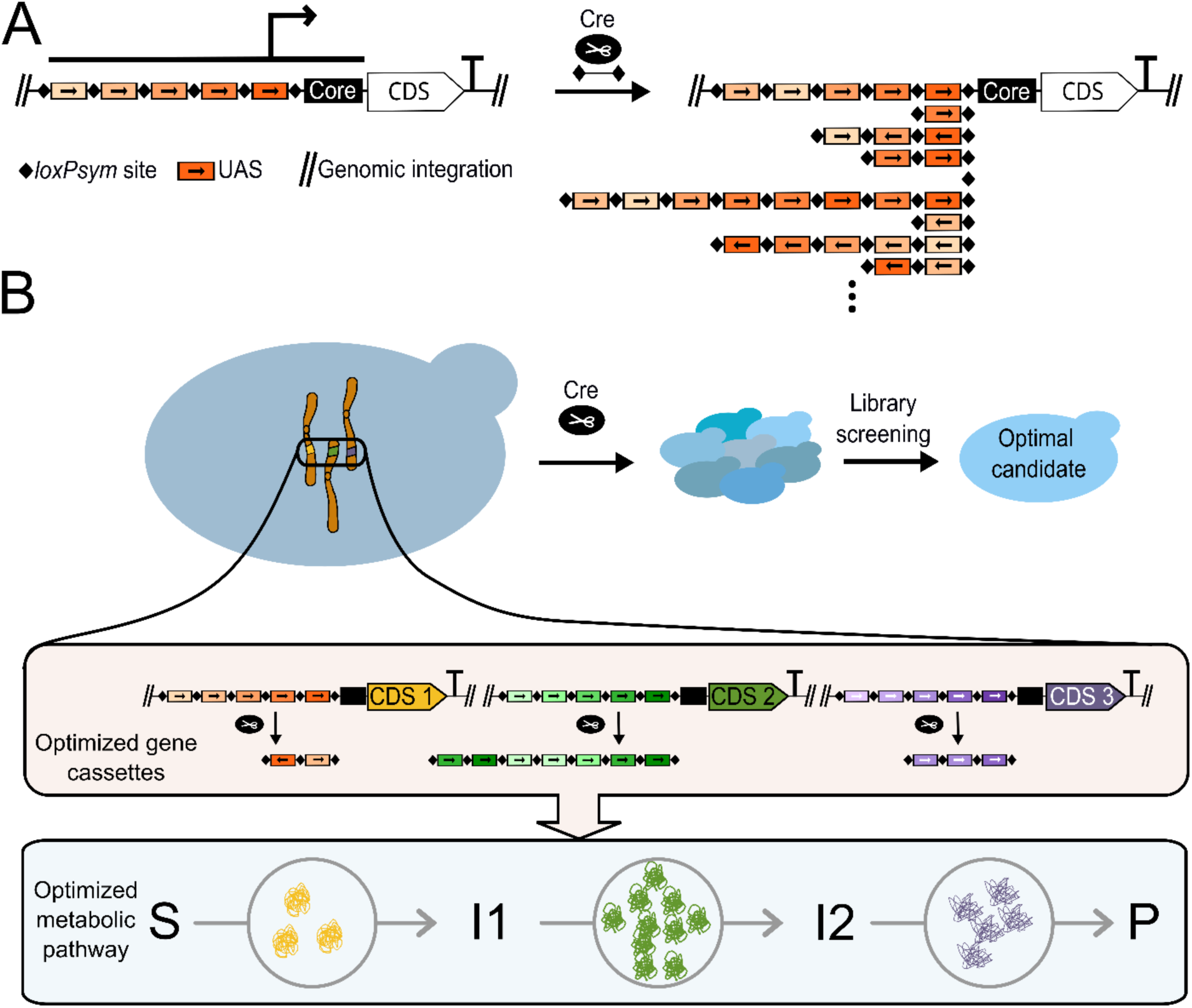
Design and application of PULSE promoters in pathway engineering. (A) Schematic overview of the design of a PULSE promoter consisting of a core promoter and five UAS elements flanked by *loxPsym* sites. Cre activity leads to deletions, inversions or insertions of individual UAS elements, resulting in a promoter library with diverse promoter configurations and promoter strengths. (B) Illustration of the PULSE tool for optimization of metabolic pathways. Genes harboring the PULSE promoters are integrated at different chromosomal locations into the genome of *S. cerevisiae*. These genes encode for metabolic enzymes that catalyze the formation of a desired bioproduct in multiple intermediate reactions. Cre-mediated recombination transforms this isogenic strain into a heterogeneous library, from which optimized candidates can be identified through screening procedures.

In the following, we present the development, validation and application of the toolkit, starting with the general design of PULSE promoter cassettes.

### FACS-based screening of randomized DNA yields active UAS elements

The design of the individual UAS elements, which served as building blocks of the PULSE promoters, was based on two central requirements: First, the expression levels of the UAS elements needed to be moderate, as they define the incremental steps in expression strength. Overly strong UAS elements would prevent a gradual increase in the range of expression levels. However, combining multiple UAS elements should result in a strong promoter cassette. Therefore, the native *PFY1* promoter, a weak and robust promoter across different conditions was used as a benchmark for the desired promoter strength of individual UAS elements (34). Second, the PULSE promoters require UAS elements of at least 82 bp to ensure sufficient distance between two *loxPsym* sites for efficient Cre-mediated recombination (35). Previously characterized native UAS elements from *S. cerevisiae* can be utilized to build promoter cassettes. However, these UAS elements vary in length but typically exceed 200 bp (36–39). For our toolkit, shorter and uniformly sized UAS elements offer significant advantages. Firstly, the overall size of the promoter is reduced, and secondly, uniformity ensures consistent distances to the core promoter across different hybrid promoter designs, simplifying the characterization of the toolkit.

To generate our synthetic UAS elements, we decided to isolate UAS elements from randomized DNA sequences. As transcription factor binding sites can be as small as 5 bp, Redden and Alper showed in 2015 that active synthetic UAS elements of a length of only 10 bp can be identified by screening randomized DNA libraries (20, 40, 41). These randomized sequences may include binding sites for native transcription factors, which can enhance the activity of a core promoter. However, adding a *loxPsym* recombination site to the promoter design was previously shown to affect the promoter activity (14, 42). Therefore, this effect needs to be considered when we screen for UAS elements.

Therefore, we utilize our specific design for the isolation of UAS elements from randomized DNA sequences. Based on the requirements for Cre-recombination, we have defined the randomized sequence part to be 48 bp. This allows the addition of 17 bp upstream and downstream of the randomized sequence to incorporate restriction enzyme binding sites, facilitating the subsequent assembly of multi-UAS promoter cassettes (Supplementary Figure 1A). Inspired by the workflow of Redden and Alper, we generated the entry vector pYTK_UP_001c as a screening platform, which holds a *loxPsym*-flanked *CcdB* sequence upstream of the previously identified minimal core 1 (20) and the reporter gene *yEGFP*. Transcription of *yEGFP* is terminated by the expression enhancing *SPG5* terminator (43). The *CcdB* gene can be used for counter selection and allows an easy exchange by the randomized DNA library in a one-step restriction and ligation reaction (Supplementary Figure 1B). This assembly yielded approximately 6,200 colonies from a single transformation in *E. coli*. The plasmid library was then isolated from *E. coli* and transferred to *S. cerevisiae*, yielding 60,000 colonies with a theoretical coverage of 10 times.

To facilitate screening for active UAS elements, a negative standard in form of a 48 bp nucleotide sequence with minimal transcriptional activity, termed nUAS, was designed. This sequence was generated using the Random DNA Generator (https://faculty.ucr.edu/~mmaduro/random.htm) with a GC content of 35%. Using the YeTFaSCo database (22) at the default setting (75% of maximum score as a minimum to be identified as a motif), a limited number of four motifs for transcription factor binding were identified (Supplementary Table 20), indicating the low activity of the nUAS sequence and therefore its suitability as a negative control for UAS activity.

To identify active UAS elements from a plasmid-based approach, the generated yeast library was sorted by FACS to isolate the top 2% of cells expressing *yEGFP*, which were then plated on CSM-URA (Figure 2A). The resulting 744 colonies were subjected to further pre-screening steps using a fluorescence plate reader, identifying 45 strains with activities higher than the nUAS, which were subsequently retested by flow cytometry and sequenced. A total of 15 unique UAS elements were isolated which were distinct from each other and the native *S. cerevisiae* sequence (Figure 2B, Supplementary Table 21). Each isolated UAS element shows stronger expression than the nUAS control, ranging between 1.2 and 3 times its expression strength. The YeTFaSCo database (22) predicts a range of 9–32 transcription factor motifs for the individual UAS elements (Supplementary Table 20), which may explain the increased activity compared to the nUAS.

**Figure 2.**
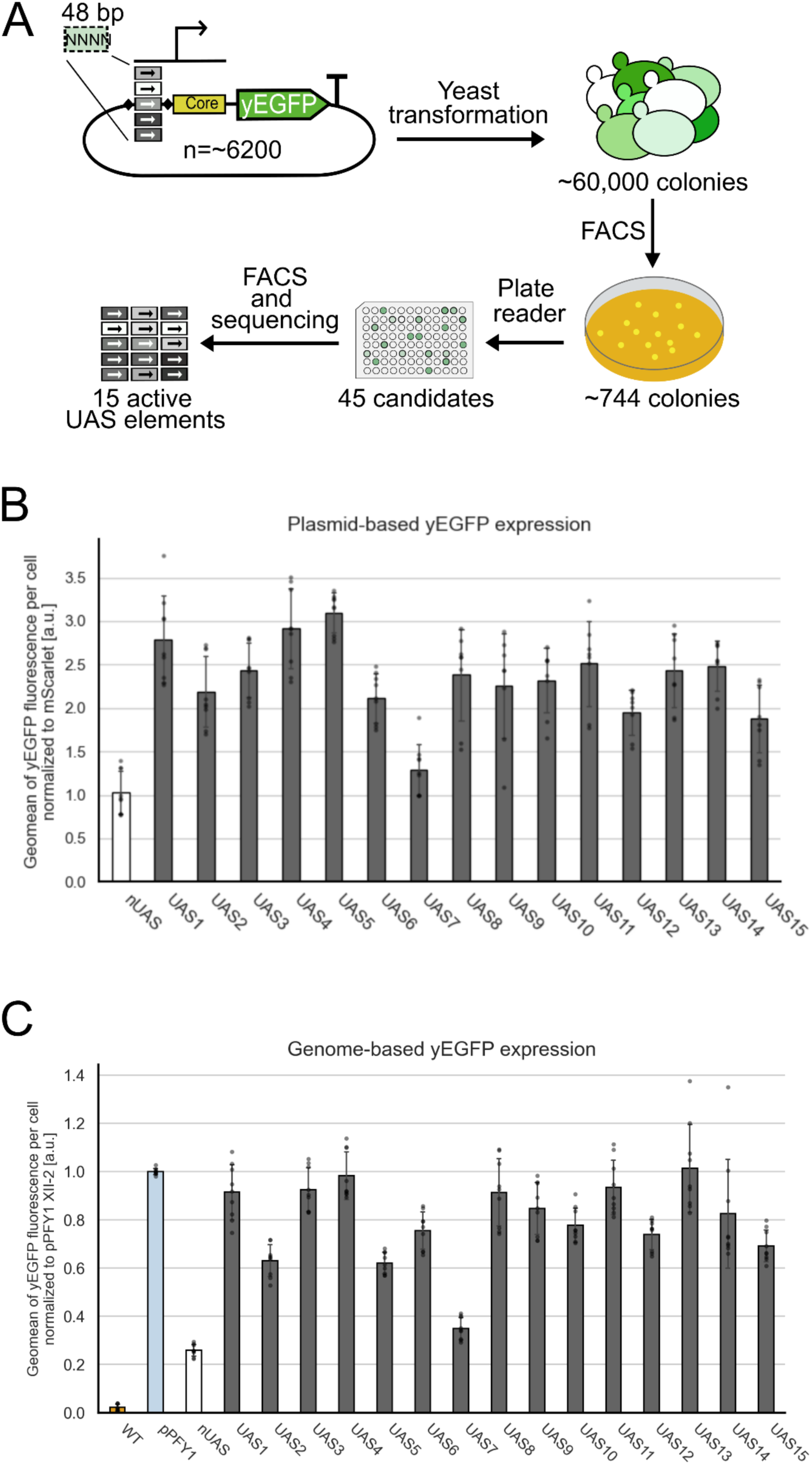
Schematic overview of the UAS isolation workflow and the performance of identified UAS elements on plasmid and chromosomal levels. (A) Randomized DNA sequences were integrated upstream of a core promoter to control *yEGFP* expression on a centromere plasmid. This plasmid library was transferred to yeast and the cells were sorted to isolate the top 2% with the highest fluorescence. The resulting colonies were retested and finally 15 active UAS elements were isolated. (B) yEGFP fluorescence mediated by the isolated UAS elements upstream of core 1 on a centromere plasmid (gray) compared to the nUAS element with only minimal activity (white). Each isolated UAS element generates a higher expression level than the nUAS control. yEGFP fluorescence was measured via flow cytometry in the exponential phase, normalized to mScarlet fluorescence (expressed by the *ADH1* promoter on the same plasmid). (C) yEGFP fluorescence mediated by the identified UAS elements upstream of core 1 compared to the WT strain (orange), native *PYF1* promoter (light blue) and the nUAS promoter (white), genome-integrated into the XII-2 locus. Each isolated UAS element (gray) shows stronger expression than the nUAS control. yEGFP fluorescence was measured via flow cytometry in the exponential phase, normalized to the expression strength of the *PFY1* promoter. For both, (B) and (C), measurements were performed in three biological replicates, each with three technical replicates (indicated as gray dots). Error bars represent the standard deviation across all replicates. a.u., arbitrary units.

To verify that the UAS elements identified in the plasmid-based screening approach exhibit consistent expression levels at the intended genomic integration site, the 15 UAS elements were genome-integrated, and their activity analyzed using flow cytometry (Figure 2C). The nUAS element and the native *PFY1* promoter as the intended benchmark were used for comparison. The single UAS hybrid promoters cover a range of approximately 40–100% of the weak native *PFY1* promoter, while the nUAS accounts for only 26%. The dataset closely aligns with the patterns observed in the episomal dataset, except for UAS5, which shifted from being the strongest UAS element in the episomal dataset to one of the weakest in the genome-integrated dataset (Figure 2B and 2C). Since we aimed for UAS element activities comparable to the PFY1 promoter, we excluded UAS2, UAS5 and UAS7, which showed only 35–63% of *PFY1* activity. However, this data demonstrates that pre-screening promoter activities at the episomal level was an effective approach for identifying UAS elements that function well in a chromosomal context.

### Combination of single UAS elements yields strong multi-UAS promoters

The assembly of multiple UAS elements is supported by a cloning strategy enabling the combination of three UAS elements using a specific restriction pattern that allows each UAS element to be assigned to a specific position in the resulting hybrid promoter (Supplementary Figure 1B). Two UAS elements can be added simultaneously to an existing promoter cassette, with one placed at the first position and the other at the last position, allowing for the creation of larger promoter cassettes containing three, five, seven or more UAS elements. The final hybrid promoter cassettes are flanked by homology regions of the yeast chromosomal locus XII-2 to enable genomic integration of the gene cassettes using the EasyClone-MarkerFree toolkit (19).

Ideal PULSE promoters should achieve high expression levels similar to strong native yeast promoters while additionally providing a smooth gradient of expression ranging from weak to strong expression levels after Cre recombination. To determine the optimal number of UAS elements for the starting promoter cassettes, the newly identified UAS elements were assembled into hybrid promoter cassettes containing 3, 5, 7, 9, 11 and 13 UAS elements, respectively. These cassettes were integrated into the XII-2 locus of the yeast genome, and *yEGFP* expression levels were measured using flow cytometry (Figure 3A, gray bars). The strengths of the hybrid promoters increased from about the level of the *PFY1* promoter with one UAS element to the level of the *ZEO1* promoter with five UAS elements to almost the strength of the *TEF2* promoter using nine to thirteen UAS elements.

**Figure 3.**
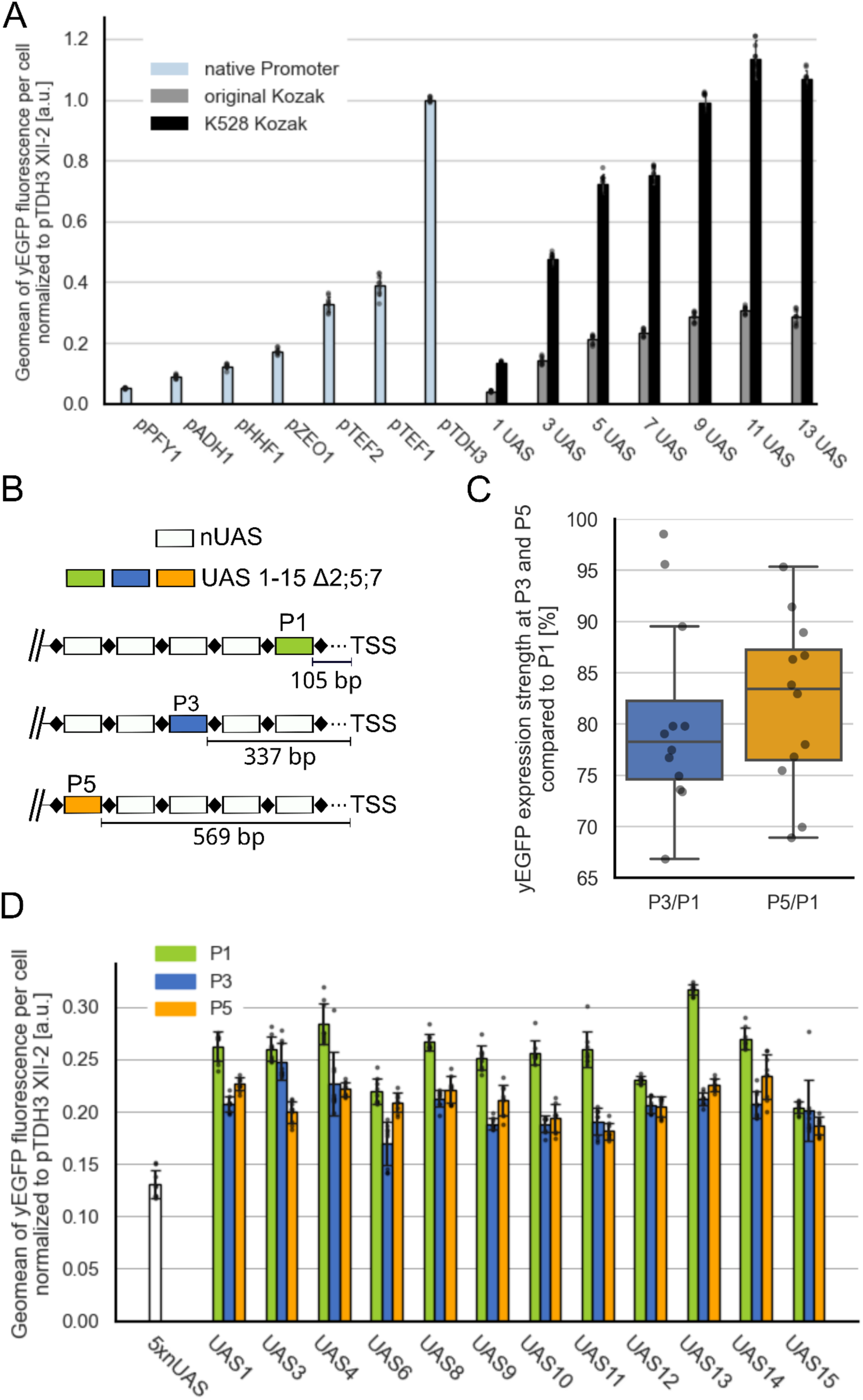
Design and testing of multi-UAS promoters. (A) yEGFP fluorescence regulated by native yeast promoters (light blue) or hybrid promoters with an increasing number of UAS elements. Gene cassettes are genome-integrated into the XII-2 locus. Hereby, the Kozak sequence is either the original (gray) or the optimized K528 sequence (black). A high number of UAS elements in combination with the K528 Kozak sequence results in the strongest promoters. (B) Schematic overview of 5xUAS promoters, in which single UAS elements are tested at three distinct positions upstream of the core promoter. The remaining positions are filled with the nUAS element. (C) Boxplot analysis of the overall activities of the UAS elements at P3 and P5 compared to P1. Around 80% activity is retained per position. (D) yEGFP fluorescence regulated by the hybrid promoters with UAS elements at either P1, 3 or 5 (XII-2 locus). All tested UAS elements showed the highest activity at P1 but also remained highly active at P3 and P5. For both, (A) and (D), yEGFP fluorescence was measured via flow cytometry in the exponential phase for three biological replicates, each with three technical replicates (indicated for each bar as gray dots). yEGFP fluorescence is normalized to the expression strength of the *TDH3* promoter in the XII-2 locus. Error bars represent the standard deviation across all replicates. a.u., arbitrary units.

To increase the overall expression output from these promoter cassettes, we used the K528 Kozak sequence, an optimized Kozak sequence identified for the core 1 promoter by Xu et al. (2021) that improves protein synthesis (Figure 3A, black bars) (21). With this, we achieved promoter activity levels that matched (in case of the 9xUAS promoter) or even surpassed (in case of the 11x and 13xUAS promoters) the expression level of the *TDH3* promoter, the strongest native constitutive promoter in *S. cerevisiae*. In both scenarios, with and without the Kozak variant K528, the additive effect of the UAS elements saturates at eleven UAS elements, with the last UAS element positioned 1,265 bp upstream the transcription start site (TSS).

While the strains harboring promoter cassettes consisting of one to seven UAS elements exhibited growth behavior comparable to that of the BY4742 wildtype strain, incorporating nine to 13 UAS elements resulted in a minor extension of the lag-phase (Supplementary Figure 2A). Nevertheless, their growth rate in the exponential growth phase is comparable to the wild type strain (Supplementary Table 22). This effect was observed for both versions of the gene expression cassettes, with and without K528, suggesting that the promoter’s strong expression strength is unlikely to be the cause. To rule out the possibility that the high number of *loxPsym* sites may form secondary structures that interfere with replication and impair cellular growth, we performed two tests:

First, each *loxPsym* site in the 3x to 11xUAS cassette was replaced with a unique, evenly sized randomized sequence with minimal predicted transcriptional activity (22) (Supplementary Figure 2B). Second, we replaced the active UAS elements with three to eleven copies of the nUAS element (Supplementary Figure 2B). Since none of these strains exhibited an extended lag-phase, we can exclude the hypothesis that inhibitory secondary structures formed by the high number of *loxPsym* sites are responsible for the observed growth effects.

Due to our limited number of twelve UAS elements, the already high expression levels of the 5xUAS promoters and the theoretical capability of the tool to generate larger and stronger promoters via Cre-mediated recombination, we decided to use and further characterize 5xUAS promoters for the tool development.

The additional effect on promoter activity from increasing the number of UAS elements suggests that our UAS elements, initially screened and identified at position one (105 bp from the TSS), remain active even at greater distances from the TSS. To examine this positioning effect, we conducted two additional experiments. First, constructs were designed in which the isolated UAS elements were tested within a 5xUAS promoter cassette at positions one, three (337 bp upstream of the TSS) and five (569 bp upstream of the TSS) (Figure 3B). The remaining positions were filled with the nUAS element. Flow cytometric analysis of yEGFP fluorescence revealed that each UAS element remained functional at positions three and position five but never exceeded the strength observed at position one (Figure 3C and 3D). On average, 80% of the expression level achieved at position one was retained at position three and five.

Second, to explore the impact of different arrangements of UAS elements on promoter strength, we tested all six possible combinations of the UAS elements UAS1, UAS3 and UAS13 in a 3xUAS configuration (Supplementary Figure 3). The expression levels of all six variants were similar, and no significant difference could be detected (ANOVA *P* = 0.2644). These findings demonstrate that the combination of isolated UAS elements has a positive cumulative effect, regardless of their order.

Our results seem reasonable compared to the natural structure of yeast promoters, where *cis*-regulatory elements are mainly located between 100 and 500 bp upstream of the TSS, but are also found in distances up to 1,400 bp (44, 45). Further support for our results comes from a 2007 study by Dobi and Winston (46) which analyzed the inducible GAL1 UAS element positioned 300-800 bp from the TATA box. The results indicated that promoters with UAS-TATA distances of up to 574 bp remain active, while those from 690 bp and beyond are inactive in one specific experimental setup. While distances conferring transcriptional activation slightly change with the genomic locus of integration and overall experimental setup, similar results were also achieved elsewhere (47, 48). It should be noted that this experiment utilized an inducible UAS element. It has been demonstrated that in native promoters, the positioning of UAS elements within the promoter shows substantial differences between inducible and constitutive promoters (49). Besides the positioning effect of UAS elements, previous studies also reported the cumulative effect of synthetic (20) and native UAS elements (36) on the promoter strength in hybrid promoter assemblies. In conclusion, it should be emphasized that transcriptional regulation is a complex interplay of various factors, including the genetic locus, the length of UAS elements and the interaction with other regulatory elements. Therefore, it is essential to test experimentally whether particular UAS elements are suitable for constructing larger hybrid promoters in the specific design.

Based on the insights gained from the experiments described so far, we designed and constructed five PULSE promoters, designated 5xUAS_A–E, to enable precise control of multi-enzymatic pathways (Figure 4A). Each of the twelve UAS elements was incorporated at least once across the construction of the five promoter cassettes. Additionally, each promoter cassette included at least three of the strongest UAS elements i.e. UAS1, UAS3, UAS4, UAS8, UAS11 and UAS13. The PULSE promoters were cloned, integrated into the XII-2 locus and yEGFP levels were analyzed. All PULSE promoters achieved similar levels of approximately 70% of the expression strength of the *TDH3* promoter (Figure 4B).

**Figure 4.**
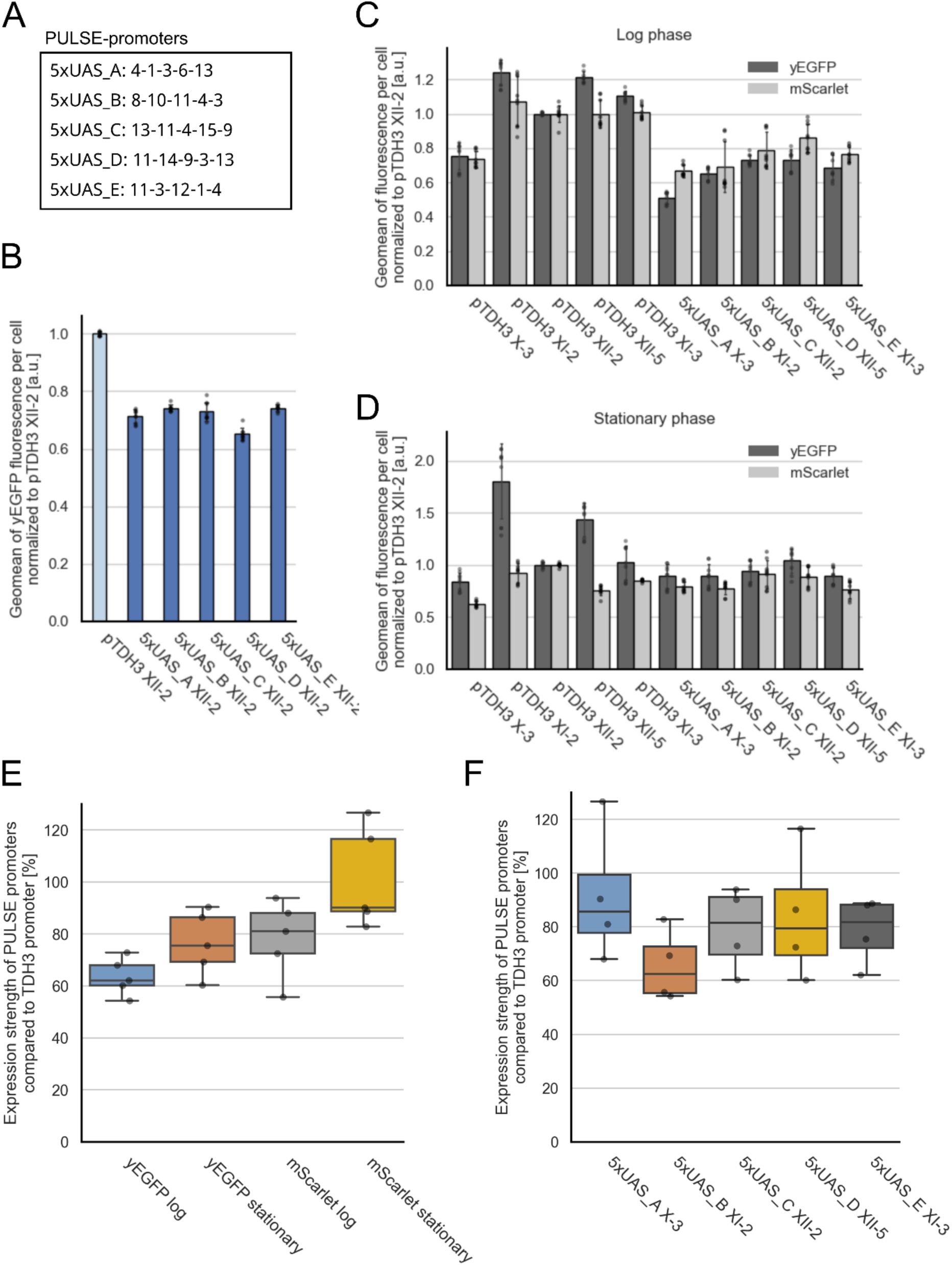
Design and testing of PULSE promoters. (A) UAS architecture of the five PULSE promoters 5xUAS_A–E. (B) yEGFP fluorescence regulated by the PULSE promoters (blue) in comparison to the *TDH3* promoter (light blue) in the XII-2 locus. All PULSE promoters reach approximately 70% expression strength of the *TDH3* promoter. yEGFP fluorescence was measured via flow cytometry in the exponential phase for three biological replicates, each with three technical replicates (indicated for each bar as gray dots). yEGFP fluorescence is normalized to the expression strength of the *TDH3* promoter in the XII-2 locus. Error bars represent the standard deviation across all replicates. a.u., arbitrary units. (C) yEGFP and mScarlet fluorescence regulated by the PULSE promoters in the log phase in comparison to the *TDH3* promoter in different genomic loci. All PULSE promoters reach strong expression levels close to that of the *TDH3* promoter. (D) yEGFP and mScarlet fluorescence regulated by the PULSE promoters in the stationary phase in comparison to the *TDH3* promoter in different genomic loci. All PULSE promoters reach strong expression levels close to the *TDH3* promoter. For (C) and (D) both, yEGFP and mScarlet fluorescence were measured via flow cytometry in the exponential phase for three biological replicates, each with three technical replicates (indicated for each bar as gray dots). yEGFP and mScarlet fluorescence are normalized to the expression strength of the *TDH3* promoter in the XII-2 locus. Error bars represent the standard deviation across all replicates. a.u., arbitrary units. (E) Boxplot analysis of the performance of the PULSE promoters 5xUAS_A–E compared to the *TDH3* promoter in the context of growth stage and CDS. The mean fluorescence of each PULSE promoter per condition was divided by the mean expression of the *TDH3* promoter in the same locus and under the same condition. Each boxplot shows the data of the five different PULSE promoters per condition. PULSE promoters reach higher expression levels in the stationary phase and furthermore higher expression levels were reached expressing *mScarlet* compared to *yEGFP*. (F) Boxplot analysis of the performance of each PULSE promoter compared to the *TDH3* promoter in the context of their genomic loci. The mean fluorescence of each PULSE promoter per condition was divided by the mean expression of the *TDH3* promoter in the same locus under the same condition. Each boxplot shows the data of individual PULSE promoters under different conditions (growth phase and CDS). 5xUAS_C–E achieve approximately 80% of TDH3 expression levels, while 5xUAS_B performs slightly worse and 5xUAS_A slightly better

To prevent unintended Cre-mediated recombination between *loxPsym* sites from different promoter cassettes, which could lead to the deletion of pathway genes in later pathway optimization experiments, the PULSE promoters will be integrated into separate genomic loci. This strategy ensures that such recombination events result in lethal strains, preventing the survival of undesirable variants. To investigate the performance of our final PULSE promoters, we studied the effect of multiple factors: First, the effect of the genomic background was investigated since all promoters have been solely tested in the XII-2 locus so far. Second, the influence of the CDS was tested, since the PULSE promoters will be used to regulate expression of different metabolic enzymes and third, the effect of the cellular growth phase on the overall expression levels of our PULSE promoters, since bioproduction usually requires growth for multiple days. Therefore, the expression levels of the PULSE promoters were tested in their designated genomic locus using yEGFP and mScarlet as reporters across the logarithmic growth phase (Figure 4C) and the stationary growth phase (Figure 4D). As a comparison, gene cassettes regulated by the *TDH3* promoter were inserted in each locus and tested accordingly. The flow cytometric data was normalized to the *TDH3* promoter in the *XII-2* locus for each condition (fluorescent protein and cellular growth phase). In both growth phases the PULSE promoters reach strong expression levels close to the *TDH3* promoter regulating both, *yEGFP* and *mScarlet*. However, small differences in expression based on the genomic loci can be observed. Gene expression in the X-3 locus turned out to be the weakest in all yEGFP constructs as well as for the *TDH3* promoter regulating *mScarlet*. Next, we compared the expression strength of each PULSE promoter relative to the *TDH3* promoter expression in the same locus throughout the different conditions as a standard for strong expression (Figure 4E). Relative to the *TDH3* promoter, PULSE promoters regulating *yEGFP* and *mScarlet* show stronger expression levels in the stationary phase compared to the logarithmic growth phase. This might be due to the fact that the *TDH3* promoter is less active in the stationary phase in CSM medium, whereas our PULSE promoters remain more stable across the two growth phases (50). Furthermore, the relative expression level of PULSE promoters when regulating *mScarlet* is higher compared to *yEGFP*.

When analyzing individual PULSE promoters in the same genomic locus, independent of the different conditions (fluorescent protein and growth phase), 5xUAS_C–E achieve approximately 80% of *TDH3* expression levels, while 5xUAS_B performs slightly worse and 5xUAS_A slightly better (Figure 4F). These dataset demonstrates that in addition to the effect of the promoter on the overall expression level, which has a key influence, the context of the CDS, the growth phase of the cell culture as well as the genomic integration site should be considered as well (19, 51, 52). Taking these high expression levels in all the tested conditions into account, the overall performance of the PULSE promoters was considered sufficient to fine-tune promoter levels of metabolic pathway enzymes.

To further enhance expression levels, if needed, there are multiple options: First, the FACS-based screening procedure of randomized DNA libraries could be repeated to find UAS elements with even higher activity. Second, even larger hybrid promoters consisting of more than 5 UAS elements can be used and lastly, genomic integrations sites can be reconsidered.

### Recombination of multi-UAS promoter cassettes generates a broad spectrum of expression levels

After establishing promising PULSE promoters, the optimal experimental conditions for Cre-recombination of UAS elements were determined, allowing us to generate a promoter library with a wide range of expression levels. The red light-inducible Cre recombinase L-SCRaMbLE was chosen to induce the recombination (53). This system is built from a split Cre recombinase which is fused to the photoreceptor PhyB and the interacting factor PIF3 from *Arabidopsis thaliana.* Upon red light induction (λ = 660 nm), PhyB and PIF3 dimerize and aid the formation of a functional Cre recombinase in the presence of the chromophore phycocyanobilin (PCB). Illumination by far-red light (λ = 740 nm) leads to dissociation of PhyB and PIF3 and therefore the Cre complex. The Light-Cre1 system (plasmid pLH_Scr15) was chosen, where only the PIF3-CreC fragment contained a nuclear localization signal to spatially separate the halves of the split protein and minimize unintended Cre activity before light induction. To define the most efficient induction time for the Cre recombinase, cells holding the 5xUAS_C promoter regulating *yEGFP* expression were illuminated with red light pulses for one to four hours and fluorescence levels of cells after recombination were analyzed via flow cytometry. Longer periods of light-induction led to an enrichment of lower *yEGFP* expressing cells in the library (Figure 5A). A light induction period of two hours was selected as the optimal induction time, as it generates a library that includes strains with weaker promoters comparable to the *PFY1* promoter while also retaining cells with stronger promoters near the strength of the *TDH3* promoter – an important prerequisite for the development of our promoter engineering tool. Recombination of larger UAS promoters (7x, 9x and 11xUAS) shows similar effects, while an induction period of three to four hours may be more appropriate for the 9x and 11xUAS promoters (Supplementary Figure 4A).

**Figure 5.**
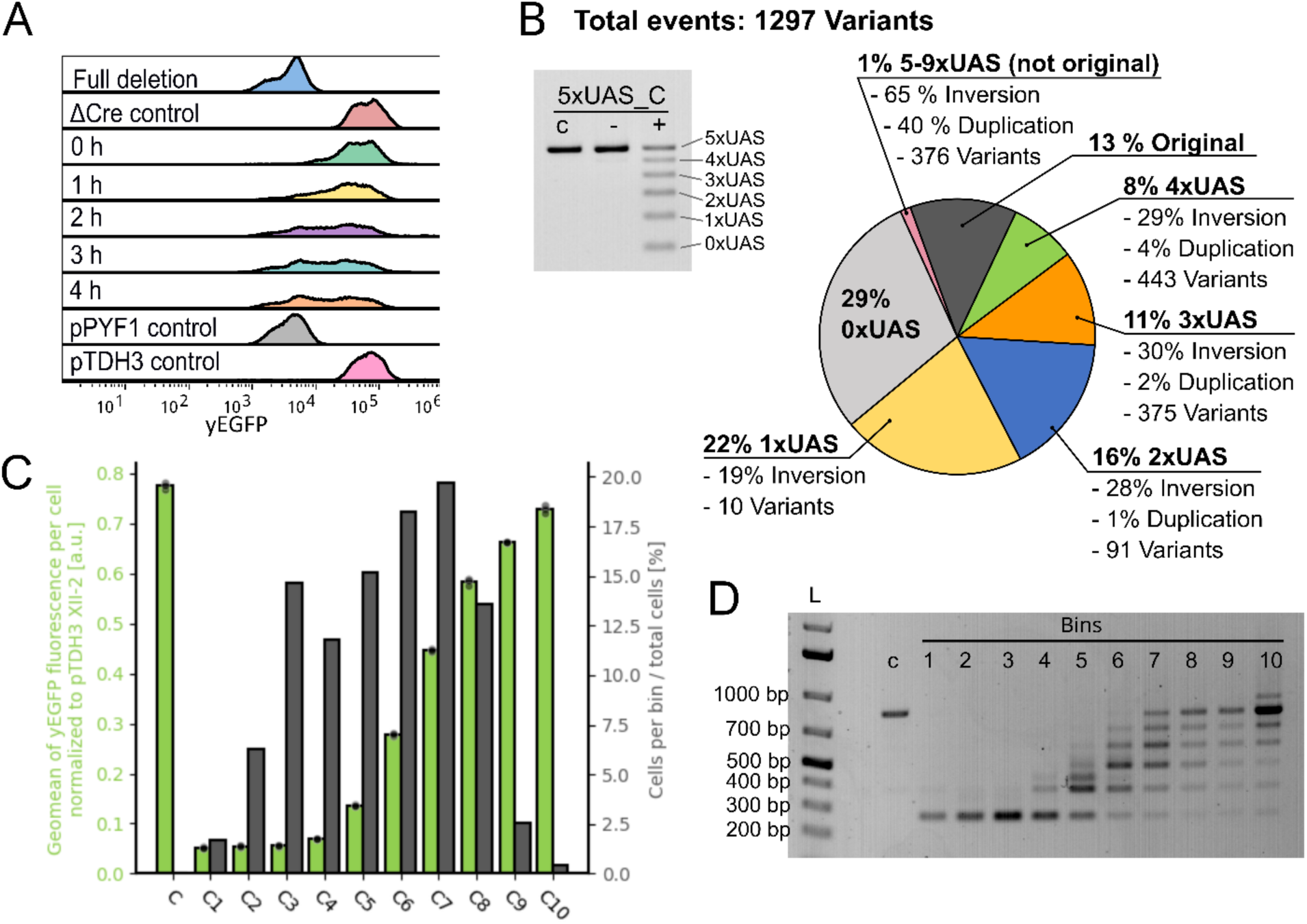
Characterization of the recombination events of the 5xUAS_C promoter. (A) Effect of different times of red-light induction on promoter strength. yEGFP fluorescence obtained from the Cre-recombined 5xUAS_C promoter library, compared to a promoter without UAS elements, the 5xUAS_C promoter in a strain without the Cre expression system and native promoter controls are shown. Longer periods of light induction led to an accumulation of cells expressing less yEGFP in the library. yEGFP fluorescence was measured via flow cytometry in the exponential phase. (B) Analysis of promoter recombination via agarose gel electrophoresis and Nanopore sequencing. PCR-amplified promoters of strains harboring the 5xUAS_C promoter, grown with red light treatment (+), grown without red light treatment (-) and of an untreated colony as a control (c), analyzed on a 2% agarose gel. The red light treated sample shows six different amplicon sizes accounting for distinct deletion events, which were further analyzed via Nanopore sequencing and summarized in a pie chart. The recombined promoters are pooled according to their inherent number of UAS elements. In addition, the proportion of promoters with inversions and duplications, as well as the total amount of different promoter architectures is given for each category. The data derives from a single run without replicates. (C) yEGFP fluorescence obtained from Cre-recombined promoters derived from the 5xUAS_C promoter library. According to the yEGFP fluorescence intensities, the library was sorted in ten uniform bins and remeasured. Shown is the mean *yEGFP* expression of cells per bin (green) as well as the distribution of cells in the library across these ten bins (gray). yEGFP fluorescence was measured via flow cytometry in the exponential phase for one biological replicate with three technical replicates (indicated for each bar as gray dots). yEGFP fluorescence is normalized to the expression strength of the *TDH3* promoter. Error bars represent the standard deviation across all replicates. a.u., arbitrary units. (D) Analysis of the PCR-amplified promoter cassettes from each bin on a 2% agarose gel, compared to the unrecombined 5xUAS_C promoter (c). While bins 1–3 mainly contain fully deleted promoter variants, a gradual increase in promoter size can be detected for the following bins.

As we decided to focus on 5xUAS promoter cassettes for further tool development, the outcome after 2 h recombination of the PULSE promoter 5xUAS_A–E was analyzed in more detail by a combination of FACS sorting and Oxford Nanopore sequencing as well as promoter size analysis by agarose gel electrophoresis: First, the promoter regions were PCR-amplified from the genome and analyzed on an agarose gel (Figure 5B for 5xUAS_C and Supplementary Figure 4B for 5xUAS_A–E). Uninduced samples show the characteristic amplicon size of a 5xUAS promoter demonstrating the tightness of our non-induced system, whereas the induced samples show six different amplicon sizes resulting from different numbers of UAS elements within the promoter. The amplicons were further analyzed by Oxford Nanopore sequencing, which revealed many different recombination events for the five promoters (Figure 5B and Supplementary Tables 16 and 23 for 5xUAS_C, and Supplementary Figure 4C and Supplementary Tables 14–15, 17–18 and 23 for 5xUAS_A, B, D, E). In all five strains, deletions were the most observed outcome. Among these, 4xUAS promoter cassettes represented the least frequent deletion outcome, accounting for approximately 4–8% of the total reads. The frequency of deletions increased with each subsequent loss of UAS elements, with complete deletion of all UAS elements being the most frequent outcome, occurring in 23–42% of the cells. Additionally, 13–19% of the cells retained the original promoter cassette. About 11–16% of the analyzed amplicons contained inversions and ∼1% contained duplications. De Boer et al. (2020) found that in transcriptional regulation, orientation of single transcription factor motifs do not show a clear relationship with their activity, indicating that inverted UAS elements might have an unpredictable effect on the overall expression strength of the hybrid promoters (54). Large promoter cassettes containing five UAS elements, different from the original configuration, or even more UAS elements were also detected, but represented only 0.6–1.5% of the total events. Overall, 561–1427 unique promoter sequences were discovered per recombined promoter library, highlighting the genomic diversity of the resulting promoter cassettes. Importantly, the distribution of individual deletion events obtained from Nanopore sequencing closely aligns with the quantification from agarose gel analysis, confirming that data processing biases did not critically affect the sequencing results (Supplementary Figure 4D).

To further demonstrate the diversity of promoter strengths obtained after light-induced recombination, each of the five yeast libraries 5xUAS_A–E, each holding a single recombined PULSE promoter, was sorted via FACS into ten uniform bins based on its fluorescence level, covering the entire population. Cells in each bin were re-cultured after sorting and subsequently re-measured by flow cytometry. For the promoter library of 5xUAS_C, an increase in promoter strength is observed from bins three to ten, spanning a range of expression levels from 5% to 73% relative to the *TDH3* promoter (Figure 5C and Supplementary Figure 4E). Approximately 40% of the cells are distributed in bins six and seven, corresponding to a promoter strength of 28–45% relative to the *TDH3* promoter. Another 40% of the cells (bins 1–5) show expression strengths of 5–14% of the *TDH3* promoter and the remaining 20% of cells (bins 8–10) account for expression strengths of 58–73% of the *TDH3* promoter. The recombined libraries of 5xUAS_D and 5xUAS_E show similar distribution patterns. In 5xUAS_A and 5xUAS_B around 60% of the cells account for expression strengths of 1–14% of the *TDH3* promoter, 15% account for approximately 27% *TDH3* expression strength and the remaining 25% account for 45–60% of *TDH3* expression strength. PCR amplification of the promoter cassettes in each bin shows that bins 1–4 mainly contain deletions of all UAS elements (Figure 5D). In the higher bins, larger promoter cassettes are observed, with bin 10 even containing 6xUAS cassettes. These datasets confirm that the recombination of the PULSE promoters 5xUAS_A–E generates highly diverse recombination events, resulting in promoters with a wide range in expression strengths that can be utilized for balancing metabolic pathways.

In applications where higher overall expression levels are required, PULSE promoter cassettes with stronger core promoters can be employed. Positioning the pTDH3 core promoter upstream of the 5xUAS_C cassette resulted in an expression level that was twice as high as that of the native *TDH3* promoter (Supplementary Figure 5A). Recombination of this promoter cassette generated a broad range of strong promoters. However, weaker promoters were underrepresented, likely because the *TDH3* core promoter alone drives expression levels higher than pPFY1 (Supplementary Figure 5B).

Based on its reliable and well-balanced performance, we selected Core 1 as the core promoter for subsequent pathway optimization experiments. Nevertheless, the *TDH3* core promoter remains a valuable component of the PULSE toolkit, offering the potential to further boost expression levels when higher output is desired. This demonstrates the flexibility of PULSE–not only in combining diverse UAS elements, but also in its compatibility with different core promoters.

### PULSE-mediated optimization of the β-carotene pathway

To showcase the capability of our toolkit to balance and optimize metabolic pathways, we applied it to enhance the production of the orange-colored pigment β-carotene. Using a pigment-producing pathway simplified the screening process by allowing visual identification of optimized candidates. To enable production of β-carotene in *S. cerevisiae*, strain β -PULSE-0 was generated where PULSE promoters were used to regulate three heterologous genes from *Xantophyllomyces dendrorhous* (*CrtE*, *CrtI*, *CrtYB-2*) and a truncated version of the native *HMG1* gene (*tHMG1*), which was previously shown to enhance the precursor production of isopentenyl pyrophosphate (IPP) and dimethylallyl pyrophosphate (DMAPP) in the mevalonate pathway (Figure 6A) (55). In β-PULSE-0, 5xUAS_A regulates *CrtE*, 5xUAS_B regulates *CrtI*, 5xUAS_C regulates *CrtYB-2* and 5xUAS_D regulates *tHMG1*.

**Figure 6.**
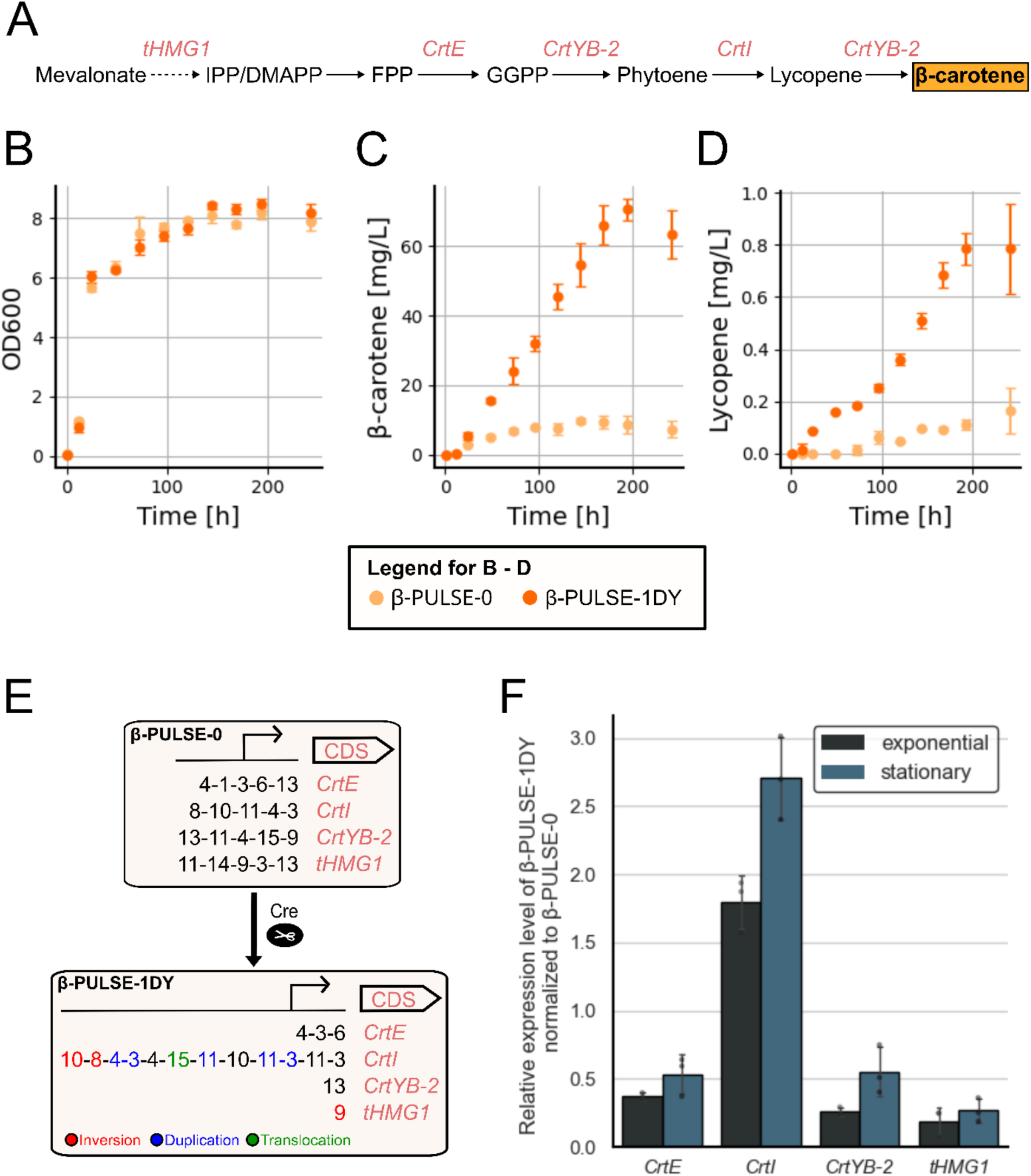
Optimization of the β-carotene biosynthesis using PULSE. (A) Schematic overview of the enzymatic steps to produce β-carotene in *S. cerevisiae*. Expression of *tHMG1* enhances the production of the precursors IPP and DMAPP in the mevalonate pathway. FPP is converted into β-carotene catalyzed by the enzymes encoded by *CrtE*, *CrtI* and *CrtYB-2*. (B-D) Analysis of β-PULSE-0 and β-PULSE-1DY cultivated in shake flasks over 240 h, focusing on (B) growth behavior, (C) β-carotene production, and (D) lycopene production. The data was generated for three biological replicates. Error bars represent the standard deviation across all replicates. (E) Illustration of the PULSE promoter architectures of the β-PULSE-0 and the optimized β-PULSE-1DY strain. (F) RT-qPCR analysis of genes regulated by PULSE promoters. Relative gene expression levels of β-PULSE-1DY were determined by using β-PULSE-0 as the reference strain for the ΔΔCT method. The data was generated for three biological replicates, each with three technical replicates. Error bars represent the standard deviation across all replicates.

In an initial test, we assessed whether multiple PULSE promoters throughout the genome cause strain instability due to the homologous recombination of repeated *loxPsym* sites, core promoters, terminators and UAS elements. Therefore, we cultivated the β-PULSE-0 strain for 100 generations (∼200 h) in a Chi.Bio reactor in the turbidostat mode (28) with dither function, which automatically diluted the culture after each generation to simplify the tracking of doublings (Supplementary Figure 6A). As a first readout, we checked the color of approximately 500 colonies, as changes in expression levels of β-carotene pathway genes typically alter pigmentation (8). All re-streaked cells retained the same color phenotype after cultivation for 100 generations (Supplementary Figure 6B). Therefore, the PULSE promoters seemed to remain unchanged. Next, we amplified each PULSE promoter cassette before and after cultivation for 100 generations and analyzed them by Oxford Nanopore sequencing (Supplementary Figure 6C). Across all PULSE promoters, recombination events after 100 generations of cultivation increased by no more than 0.36% compared to the starting culture, indicating that the strains are genomically stable in the absence of a functional Cre recombinase.

Subsequently, promoter recombination in β-PULSE-0 was induced using the red-light inducible Cre recombinase, and the resulting β-PULSE-library was plated on agar. The PULSE promoter recombination generated six distinct cell phenotypes, ranging in colors from orange to various shades of yellow to pink and white. Beta-carotene was extracted from four different colonies of each phenotype and quantified by HPLC (Supplementary Figure 7). It was determined that cells of the orange phenotype contained the highest β-carotene levels, thus seven more orange colonies were tested. The best performer, strain β-PULSE-1D, turned out to be a mixture of a yellow (β-PULSE-1DY) and an orange phenotype (β-PULSE-1DO). Surprisingly, after isolation and HPLC analysis of β-PULSE-1DY and β-PULSE-1D0, the former turned out to produce more β-carotene and was used for further characterization.

While both β-PULSE-0 and β-PULSE-1DY showed similar growth behavior (Figure 6B), β-PULSE-1DY produced approximately 24 mg/L β-carotene after 72 h and up to 70 mg/L after 194 h of shake flask cultivation, representing up to an eightfold increase compared to β-PULSE-0 (Figure 6C). Lycopene, the direct precursor of β-carotene, accumulated at very low levels in both strains, with yields below 1 mg/L (Figure 6D). Sequencing the PULSE promoters in β-PULSE-1DY revealed the following configurations: *CrtE* was found to be regulated by a 3xUAS promoter, *CrtI* by a 10xUAS promoter, *CrtYB-2* by a 1xUAS promoter, and *tHMG1* by a 1xUAS promoter (Figure 6E). While deletions are generally favored in *loxPsym*-based recombination events (56, 57), the *CrtI* promoter in β-PULSE-1DY showed a complex rearrangement pattern, including duplications, inversions, and even a translocation from a different genomic locus. Notably, UAS15, which was not part of the original promoter cassette, must have translocated from the 5xUAS_C promoter that regulates *CrtYB-2* at the XII-2 locus to the XI-2 locus of *CrtI*. Initial PCR-based Sanger sequencing revealed a 2xUAS cassette upstream of *CrtI* in β-PULSE-1DY. However, the corresponding transcript levels did not match this configuration. Whole genome sequencing subsequently identified a 10xUAS cassette at the same locus. The high number of repeats likely interfered with PCR amplification of this specific promoter, as polymerase skipping has been reported for other repetitive DNA sequences (58). Notably, this was the only case in which PCR-based detection failed; other multi-UAS promoter cassettes — including configurations with up to 13xUAS — were successfully amplified and verified by Sanger sequencing without issues.

To analyze gene expression levels of PULSE promoters in β-PULSE-1DY, RT-qPCR was performed for all four β-carotene pathway genes during exponential and stationary growth phases. Expression levels were compared to those in β-PULSE-0, in which each gene is regulated by a 5xUAS promoter (5xUAS_A-D), using the ΔΔCT method (Figure 6F). As expected, *CrtE*, *CrtYB-2* and *tHMG1*, which are regulated by promoters containing fewer than five UAS elements in β-PULSE-1DY, showed reduced transcript levels compared to the corresponding genes in β-PULSE-0. In contrast, *CrtI*, which is driven by a 10xUAS promoter, showed strongly increased expression, reaching 170% in exponential and 260% in stationary growth phase, relative to the 5xUAS_B-driven expression in β-PULSE-0. Overexpression of *CrtI* had been shown to enhance β-carotene biosynthesis before (55), suggesting that the 10xUAS promoter plays a critical role in the good performance of this strain. The identification of this 10xUAS promoter among a limited number of tested colonies highlights PULSE’s ability to generate exceptionally strong expression when needed. Remarkably, the high *CrtI* expression level remained stable in the stationary phase, likely explaining the continuous increase in β-carotene levels up to 200 h of cultivation.

In addition to *CrtI*, overexpression of *tHMG1* has also been proposed to enhance β-carotene biosynthesis (59, 60). However, β-PULSE-1DY utilizes a weak 1xUAS promoter for its regulation. Also, the other strong β-carotene producers (1B, 1F, 1H and 1J) utilize weak promoters consisting of one or even no UAS element to regulate *tHMG1* expression (Supplementary Figure 8). This demonstrates the capability of PULSE to not only optimize metabolic pathways but also reveal expression optimization targets.

### PULSE-mediated pathway optimization of the xylose oxidoreductase pathway

To demonstrate, that besides bioproduction also the growth on alternative carbon sources can be optimized using PULSE, we aimed to improve growth on xylose through the oxidoreductase pathway. Here as well, screening is simplified as cell growth can be used as a direct indicator for improved strains. Therefore, three heterologous genes from *Pichia stipitis* (*Xyl1*, *Xyl2* and *Xyl3*) were introduced into *S. cerevisiae* to form strain X-PULSE-0 (Figure 7A). Additionally, the native transaldolase gene *TAL1*, which was previously reported to enhance the pentose phosphate pathway, was targeted for promoter optimization to further improve xylose utilization (61, 62). In X-PULSE-0 5xUAS_A regulates *Xyl3*, 5xUAS_B regulates *TAL1*, 5xUAS_C regulates *Xyl1* and 5xUAS_D regulates *Xyl2* (Figure 7B).

**Figure 7.**
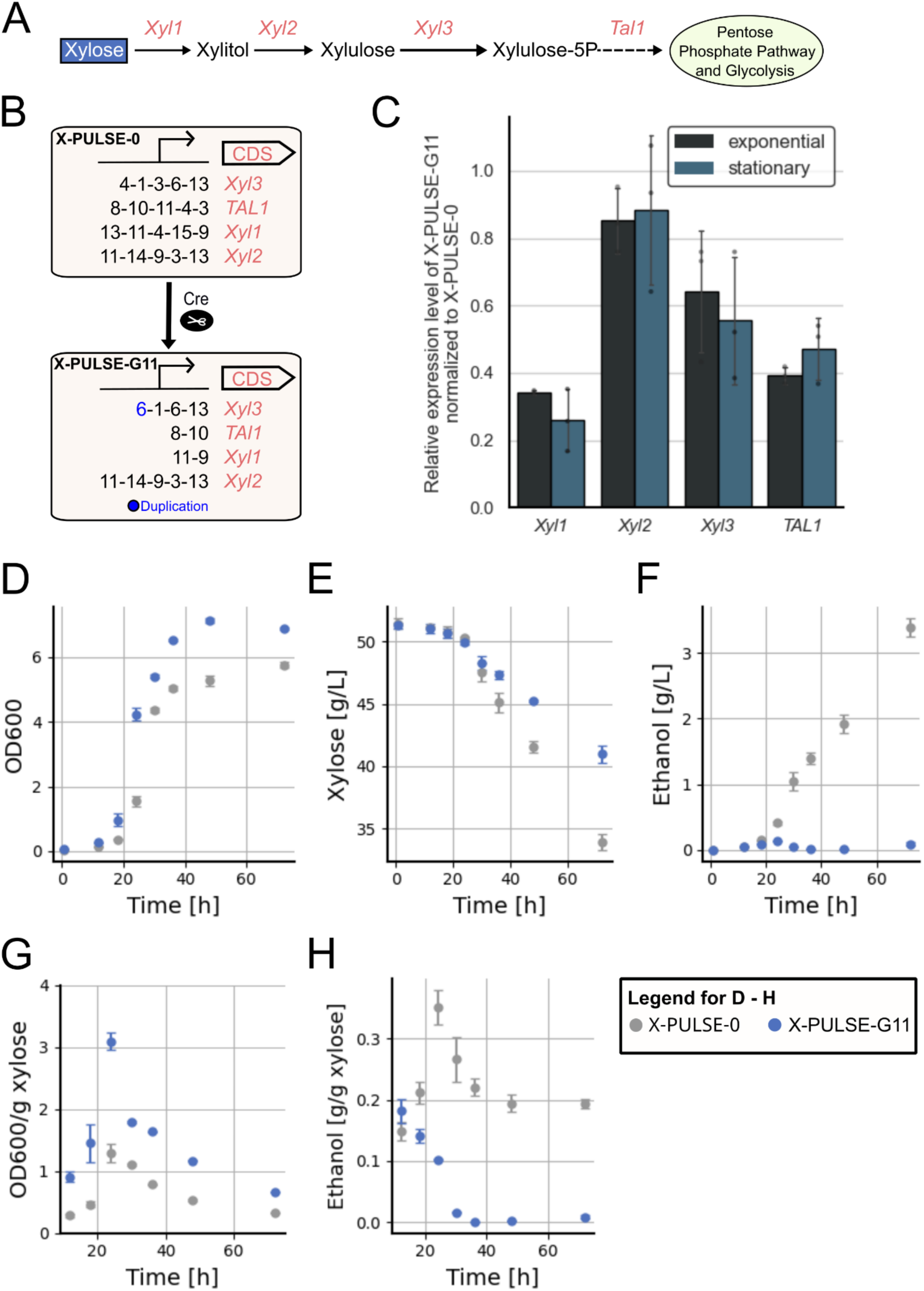
Optimization of the oxidoreductase pathway using PULSE. (A) Schematic overview of the enzymatic steps necessary for *S. cerevisiae* to utilize xylose as the sole carbon source. Xyl1-3 are necessary for xylose to enter the pentose phosphate pathway which is connected to glycolysis in *S. cerevisiae*. The *TAL1* expression level further influences the pentose phosphate pathway. (B) Illustration of the PULSE promoter architecture of the X-PULSE-0 and the optimized X-PULSE-G11 strain. (C) RT-qPCR of the genes regulated by the PULSE promoters. Relative gene expression levels of X-PULSE-G11 were determined by using X-PULSE-0 as a reference strain for the ΔΔCT method. The data was generated for three biological replicates, each with three technical replicates. Error bars represent the standard deviation across all replicates. (D-H) Analysis of X-PULSE-0 and X-PULSE-G11 cultivated in shake flasks over a period of 72 h focusing on (D) growth behavior, (E) xylose consumption, (F) ethanol production, (G) growth per consumed xylose and (H) produced ethanol per consumed xylose. The data was generated in three biological replicates. Error bars represent the standard deviation across the replicates.

Following promoter recombination of X-PULSE-0 by the red-light induced Cre recombinase, the resulting X-PULSE-library was cultured for two days in CSM liquid medium with either 10 or 50 g/L xylose as the sole carbon source to enrich faster-growing strains, before being streaked onto agar plates with 50 g/L xylose. As no obvious growth differences were visible on the plates, 47 colonies from both plates were randomly selected and tested in growth assays, evaluating both growth rate and final cell density. Based on these criteria, the four best strains were selected and further analyzed in different xylose concentrations (5, 10, 30 and 50 g/L) over a period of 82 h at multiple time points across the exponential and stationary growth phase (Supplementary Table 24). The best performing strain, X-PULSE-G11, is shown in Supplementary Figure 9 in comparison to the unrecombined strain X-PULSE-0. At a xylose concentration of 5 g/L the X-PULSE-0 strain shows faster growth. Strain X-PULSE-G11 grows equally fast compared to X-PULSE-0 at xylose concentrations of 10 g/L and furthermore reaches a higher final cell density. Remarkably, in media containing xylose concentrations of 30 and 50 g/L, X-PULSE-G11 outperforms X-PULSE-0 in the final cell density and initiates the exponential growth phase faster.

Sequencing the PULSE promoters regulating the oxidoreductase pathway genes in strain X-PULSE-G11 revealed that *Xyl1* is regulated by a 2xUAS promoter and *Xyl2* by a 5xUAS promoter, which resembles the original, non-recombined hybrid promoter. *Xyl3* is regulated by a 4xUAS promoter and *TAL1* by a 2xUAS promoter (Figure 7B).

Transcript levels of the four genes in X-PULSE-G11 were analyzed by RT-qPCR during exponential and stationary growth phases using the ΔΔCT method, with X-PULSE-0 as the reference strain (Figure 7C). Gene expression of *Xyl1* and *TAL1*, both driven by 2xUAS promoters, was downregulated to approximately 30% and 40%, respectively, compared to the expression level of the 5xUAS promoter. *Xyl3*, regulated by a 3xUAS promoter showed ∼60% relative expression, and *Xyl2* under the control of the original 5xUAS promoter exhibited a relative expression level of ∼90%. Relative expression levels were consistent across both growth phases.

For further phenotypic analysis, growth, xylose consumption, and production of xylitol and ethanol were analyzed in X-PULSE-0 and X-PULSE-G11 cultivated in baffled flasks containing 50 g/L xylose over 72 h (Figure 7D-H). X-PULSE-0 remained in the lag-phase for 9.6 ± 0.3 h and reached a maximum growth rate of 0.24±0.01 h^-1^, whereas X-PULSE-G11 exhibited a shorter lag-phase of 7.1±1.3 h and a maximum growth rate of 0.26 ± 0.02 h^-1^ (Figure 7D). X-PULSE-0 reached its highest cell density of OD_600_ of 5.8 ± 0.1 after 72 h, while X-PULSE-G11 reached a maximum OD_600_ of 7.1 ± 0.1 after only 48 h.

Unexpectedly, xylose consumption was higher in X-PULSE-0, which consumed 16 g/L xylose after 72 h, compared to 9 g/L utilized by X-PULSE-G11 in the same time period (Figure 7E). Xylitol was not detectable in either strain, indicating efficient flux through the oxidoreductase pathway and no relevant cofactor imbalance, a well-known bottleneck that often results in xylitol accumulation in *S. cerevisiae* under anaerobic growth conditions (63).

Ethanol production differed markedly: X-PULSE-0 produced 3 g/L ethanol, whereas X-PULSE-G11 produced only 0.1 g/L ethanol over 72 h (Figure 7F). X-PULSE-0 channeled xylose towards both ethanol production and biomass formation, whereas X-PULSE-G11 primarily used it for biomass synthesis (Figures 7G and H). These results suggest that, although X-PULSE-0 was cultivated under aerobic conditions, it favored fermentation over respiration due to the high sugar concentration. This is consistent with the Crabtree effect, which is normally associated with growth on glucose, but has also been observed in *S. cerevisiae* grown on high xylose concentrations (64).

While fermentation yields higher ethanol levels, it generates less ATP, a limiting resource for biomass production (65). This suggests that the downregulation of *Xyl1*, *Xyl3* and *TAL1* in X-PULSE-G11 has shifted the yeast metabolism towards respiration, explaining its improved growth performance. This could also explain why X-PULSE-G11 does not outperform X-PULSE-0 at lower xylose concentrations (e.g., 5 or 10 g/L; Supplementary Figure 9). At these concentrations, glycolytic flux in X-PULSE-0 likely remains within the respiratory capacity and does not trigger a metabolic switch to fermentation. While ethanol production in *S. cerevisiae* is industrially valuable, to yeast it is merely a by-product for cofactor regeneration (66). With its low ethanol production and efficient growth, X-PULSE-G11 can maximize carbon flux toward desired bioproducts instead of ethanol, rendering it as an interesting candidate for sustainable bioproduction from lignocellulosic biomass (67). In case improved growth at lower xylose concentrations or enhanced ethanol production is desired, future screening strategies must be adapted to include these specific criteria, as different UAS combinations may perform better under such conditions.

## Conclusion

In this study, we established an efficient workflow for the FACS-based identification of synthetic promoter elements at the plasmid level, which can be combined to construct exceptionally strong promoters. A key feature of the PULSE tool is the integration of *loxPsym*-recombination sites flanking the promoter elements. Activation of the red light-regulated Cre recombinase L-SCRaMbLE (53) enables the generation of a diverse promoter library, including hundreds of variants with deleted, inverted, translocated, or duplicated promoter elements. From a yeast library of only 6,200 variants resulting from a single transformation, we identified twelve active UAS elements with medium strengths comparable to the *PFY1* promoter, which were combined into five strong promoter cassettes and subsequently used for genomic integration and the generation of the platform strain. The promoter strength is adjusted by the number of small UAS elements in a hybrid promoter design, which can be decreased by deletions or increased by duplications. This enables theoretically unlimited promoter combinations and allows for a wide range of expression levels. Furthermore, the isolated UAS elements robustly activated not only the Core 1 promoter used in the screening procedure to identify the UAS elements, but also the native *TDH3* core promoter, enabling our hybrid promoter to double the expression level of *TDH3*–the strongest known native yeast promoter.

Using Oxford Nanopore sequencing and FACS screening, we demonstrated that recombination produces a variety of different promoter cassettes and a smooth range of expression levels. However, while the ability to generate large promoter libraries is an advantage, identifying the optimal candidate can be challenging without straightforward readouts, such as color, growth, pH, or temperature tolerance. To circumvent this challenge, we demonstrated the capability of PULSE by increasing β-carotene production eightfold and improving growth at high xylose concentrations in *S. cerevisiae*. Notably, this improvement was achieved in comparison to the parental strain carrying all genes of interest under regulation of strong promoter cassettes. Remarkably, β-PULSE-1DY produces high β-carotene levels of 70 mg/L despite being derived from a laboratory strain in which only the biosynthetic pathway genes and *tHMG1* were subjected to promoter fine-tuning (68). This is attributable to a 10xUAS promoter featuring complex rearrangements, including deletions, inversions, and even a translocation from a distant genomic region. Thus, the ability of PULSE to generate exceptionally strong promoters is demonstrated, surpassing even the strength of the *TDH3* promoter. These results underscore the potential of PULSE as a powerful tool for evolutionary experiments, where selection pressure can drive the emergence of highly active promoter variants.

Even though PULSE and the previously published GEMbLeR tool (14) share the underlying concept of using Cre recombinase to generate diverse hybrid promoter cassettes for fine-tuning gene expression *in vivo*, PULSE offers several advantages in terms of feasibility and implementation:

First, instead of relying on limited pre-characterized native promoter elements, PULSE identifies small and equally sized UAS elements through a simple screening process using a small library. This screening process could easily be repeated to identify more UAS elements and could even be performed in other non-model yeasts to rapidly identify active UAS elements.

Second, UAS elements identified by PULSE exhibit consistent activity regardless of their position within the promoter cassette, resulting in additive effects that simplify promoter construction and allow for very high overall expression levels. This represents a clear advantage over designs in which the promoter element proximal to the core promoter predominantly defines the expression level, thereby limiting the diversity of achievable expression profiles.

Third, instead of relying on a single repeatedly used promoter cassette, PULSE provides a set of well-characterized hybrid-promoter cassettes that differ in the number and identity of UAS elements as well as in the core promoter. This diversity of promoter sequences promotes the genomic stability of our strains by reducing the likelihood of homologous recombination between promoter cassettes.

The push to replace fossil-based processes with circular, sustainable bio-based alternatives demands reliable solutions for engineering robust and efficient production hosts capable of realizing economically viable processes. PULSE demonstrates its capability to address these key challenges in strain engineering by enabling precise and dynamic control of gene expression while avoiding cloning-intensive workflows and overcoming transformation efficiency limitations. Non-conventional yeasts with beneficial characteristics for bioeconomy are increasingly reported, but their potential remains limited by challenges in genomic engineering, particularly transformation efficiency (69, 70). With its cloning- and transformation-independent workflow and reliance on small synthetic UAS elements identified via simplified screening, PULSE offers a promising solution for expanding biotechnological applications to less-characterized yeasts.

## Supporting information

Supplementary_Tables_PULSE

## Data availability

All data supporting the findings of this study are available within the paper and its supplementary information files. All DNA sequences are available as GenBank files. All material created within this study is available from the corresponding author upon request. Nanopore data has been deposited to NCBI Sequence Read Archive database under accession code PRJNA1197365. Open source code for the program to analyze the hybrid promoter recombination events is available at https://doi.org/10.6084/m9.figshare.28203740.v2 and https://github.com/Tafazz/ScramblePJ.

## Author contributions

L.H. developed the overall strategy and supervised the work. C.R., A.T.Y. and L.H. designed the experiments and analyzed the data. C.R. and A.T.Y. conducted the experiments. C.R. and L.H. wrote the manuscript with contributions from A.T.Y.

## Acknowledgements

We thank Prof. Dr. Eckhard Boles (Goethe University Frankfurt), Prof. Dr. Andrea Braeutigam (University of Bielefeld), and Prof. Dr. Bernd Mueller-Roeber (University of Potsdam) as well as members of the Junior Research Group TAILOR for fruitful and inspiring discussions. We thank Dr. Mislav Oreb and Lara Junghans (both Goethe University Frankfurt) for the metabolite analysis of the xylose utilization strains. We thank the Department of Molecular Biology at the University of Potsdam for technical support throughout the study. The authors used OpenAI’s ChatGPT (model GPT-4o) to adjust the manuscript’s English.

## Supplementary Information

**Supplementary Figure 1.**
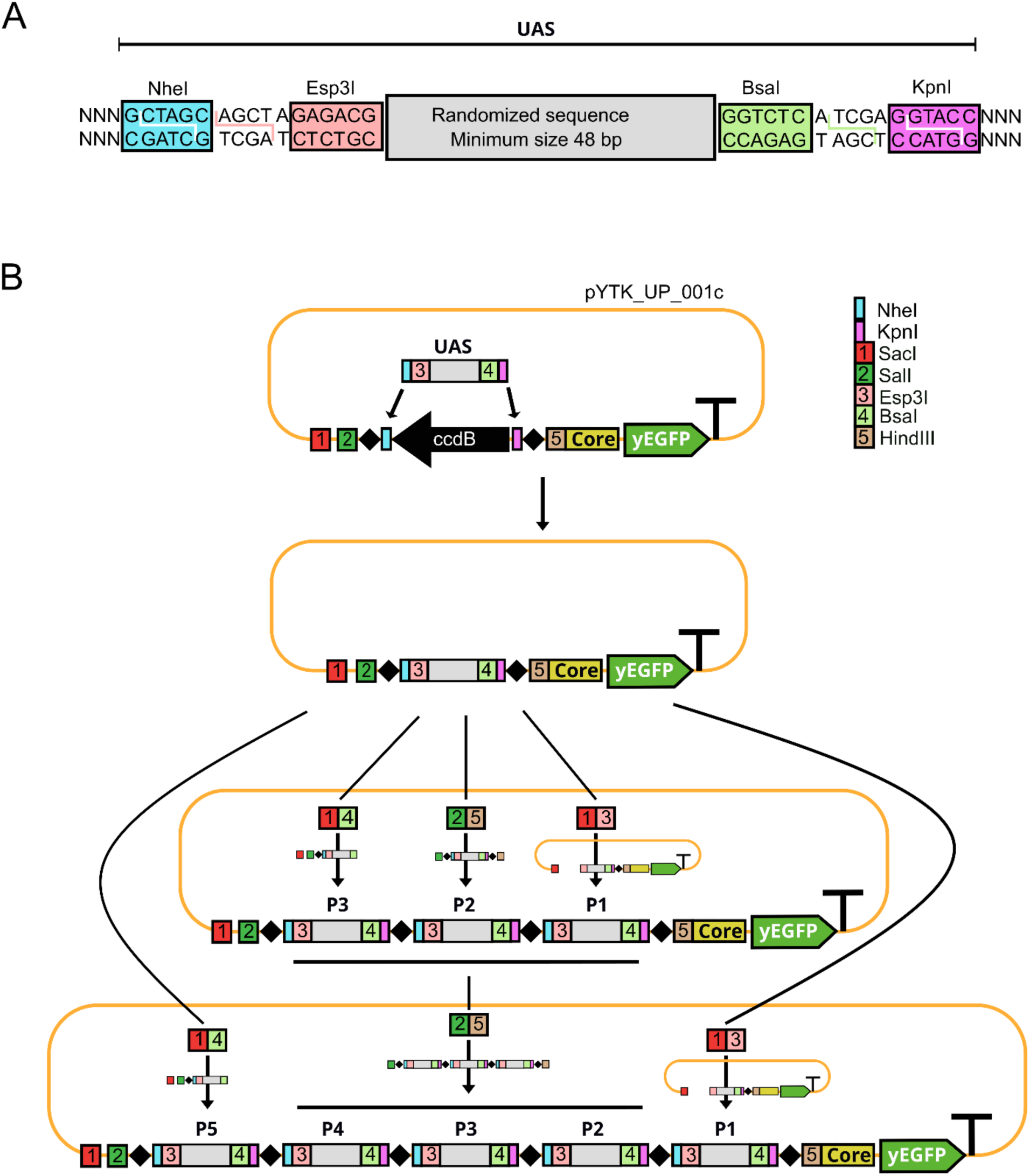
Assembly strategy of single and multi-UAS constructs. (A) Detailed depiction of the restriction sites necessary to insert single UAS elements into the entry vector pYTK_UP_001c. (B) Schematic overview of the assembly of multi-UAS constructs. First single UAS elements must be added via a restriction and ligation reaction to the entry vector. Subsequently by following specific restrictions and ligation patterns, two UAS elements can be added at a time to a hybrid promoter. This process can be repeated infinitely while always extending the promoter by two UAS elements. A detailed description can be found in the material and methods section.

**Supplementary Figure 2.**
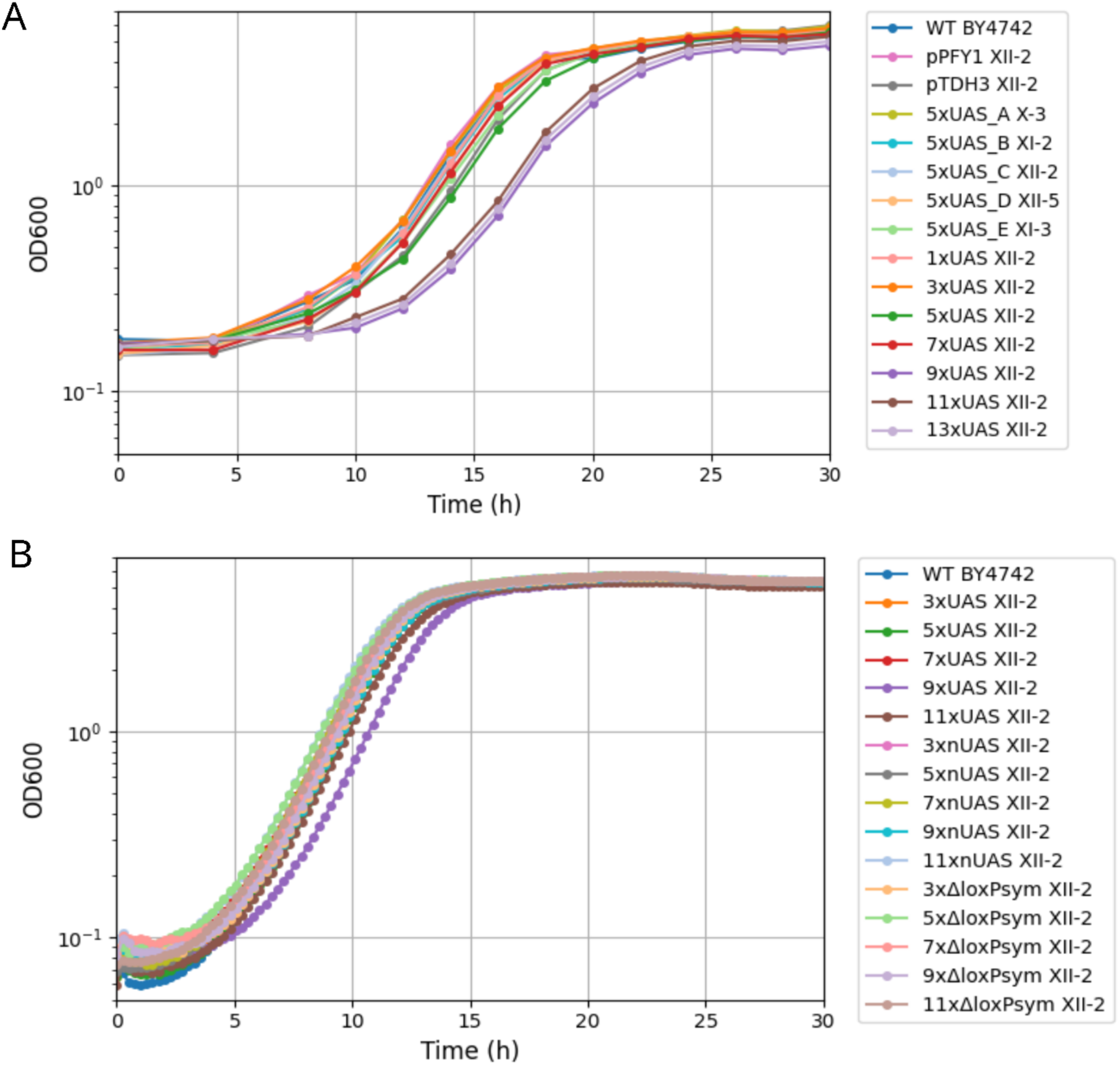
Growth curve of BY4742 with genome integrated hybrid promoters. (A) Growth of the wildtype BY4742 strain, as well as its derivates holding native and PULSE promoters regulating *yEGFP* expression were analyzed over a period of 38 h in 48-well plates at 220 rpm. The strains harboring native promoter controls (pPFY1 and pTDH3), the PULSE promoter cassette strains (A–E) and the 1x-7xUAS promoter show similar growth behavior has the wildtype strain. The strains harboring the 9x-13xUAS promoters have an extended lag-phase. Data is presented as the mean of two biological replicates with three technical replicates (B) Growth test of multi-nUAS promoters and multi-UAS promoters without *loxPsym* sites (Δ*loxPsym)* in a microplate reader. All strains show a similar growth behavior as the WT. Data is shown as the mean of three biological replicates with three technical replicates.

**Supplementary Figure 3.**
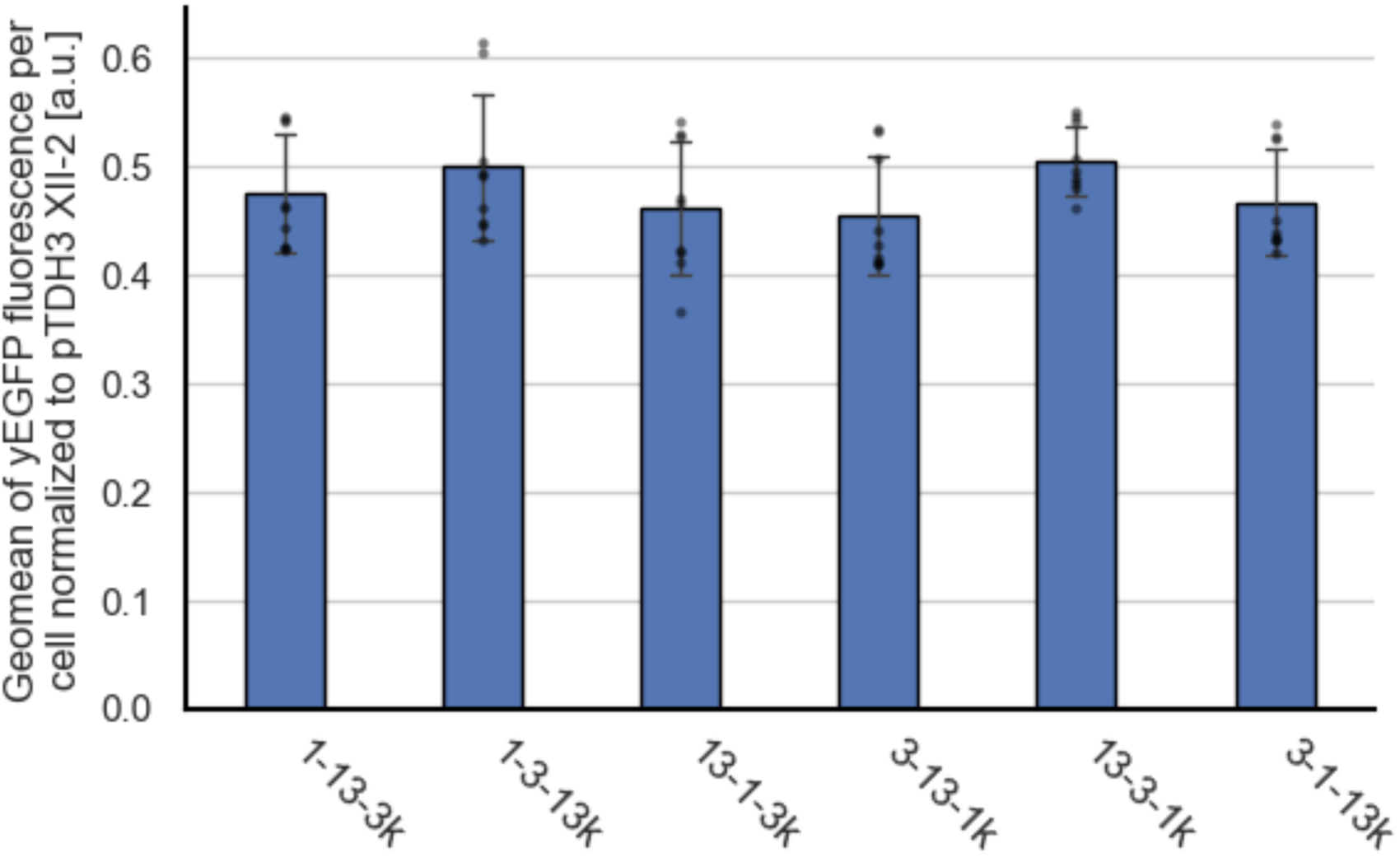
Positional effect of UAS elements in a 3xUAS promoter. yEGFP fluorescence regulated by 3xUAS promoters with all combinations of the UAS elements 1, 3 and 13 upstream of the core 1 promoter with the K528 Kozak sequence, genome-integrated into the XII-2 locus. No significant difference in expression strength can be determined. yEGFP fluorescence was measured via flow cytometry in the exponential phase for three biological replicates, each with three technical replicates (indicated for each bar as gray dots). Error bars represent the standard deviation across all replicates. a.u., arbitrary units. A one-way ANOVA revealed no significant difference in between the 3xUAS promoters (P = 0.2644).

**Supplementary Figure 4.**
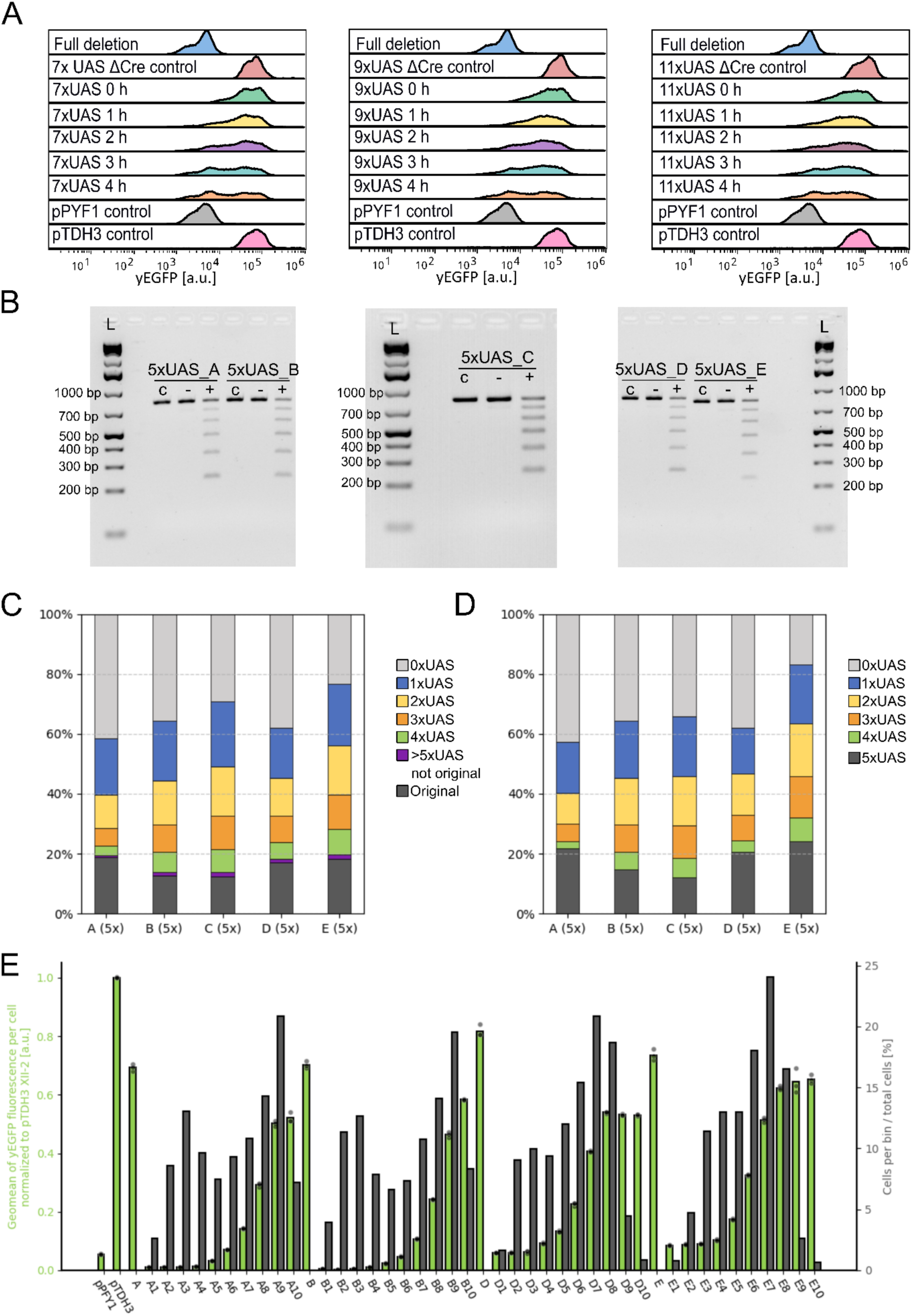
Characterization of the recombination events on different hybrid promoters. (A) Effect of different times of red-light induction on promoter strength. yEGFP fluorescence obtained from Cre-recombined 7/9/11xUAS promoter libraries compared to a promoter without UAS elements, the 7/9/11xUAS promoter in a strain without the Cre expression system and native promoter controls are shown. Longer periods of light induction led to an accumulation of cells expressing less yEGFP in the library. yEGFP fluorescence was measured via flow cytometry in the exponential phase. (B) Analysis of promoter recombination of the PULSE promoters via agarose gel electrophoresis and Nanopore sequencing. PCR-amplified promoters of strains grown with red light treatment (+), grown without red light treatment (-) and from an untreated colony as a control (c), analyzed on a 2% agarose gel. The red light treated samples show six different amplicon sizes accounting for distinct deletion events. (C) Analysis of recombined PULSE promoters via Nanopore sequencing. The recombined promoters are pooled according to their inherent number of UAS elements. The data derives from two runs without replicates (first run 5xUAS_A, B, D, E; second run 5xUAS_C, D). (D) Analysis of recombined PULSE promoters via quantification from the agarose gel picture. Each amplicon accounted for promoters with a specified number of UAS elements. The total amount of each deletion event was quantified and plotted for each PULSE promoter. The data closely aligns with the data obtained by Nanopore sequencing. (E) yEGFP fluorescence obtained from Cre-recombined promoters derived from 5xUAS_A, B, D, E. In accordance with the intensity of yEGFP fluorescence, the library was sorted in 10 uniform bins and remeasured. Shown is the mean yEGFP fluorescence of cells per bin (green) as well as the distribution of cells in the library across these 10 bins (gray). yEGFP fluorescence was measured via flow cytometry in the exponential phase for one biological replicate with three technical replicates (indicated for each bar as gray dots). yEGFP fluorescence is normalized to the expression strength of the *TDH3* promoter. Error bars represent the standard deviation across all replicates. a.u., arbitrary units.

**Supplementary Figure 5.**
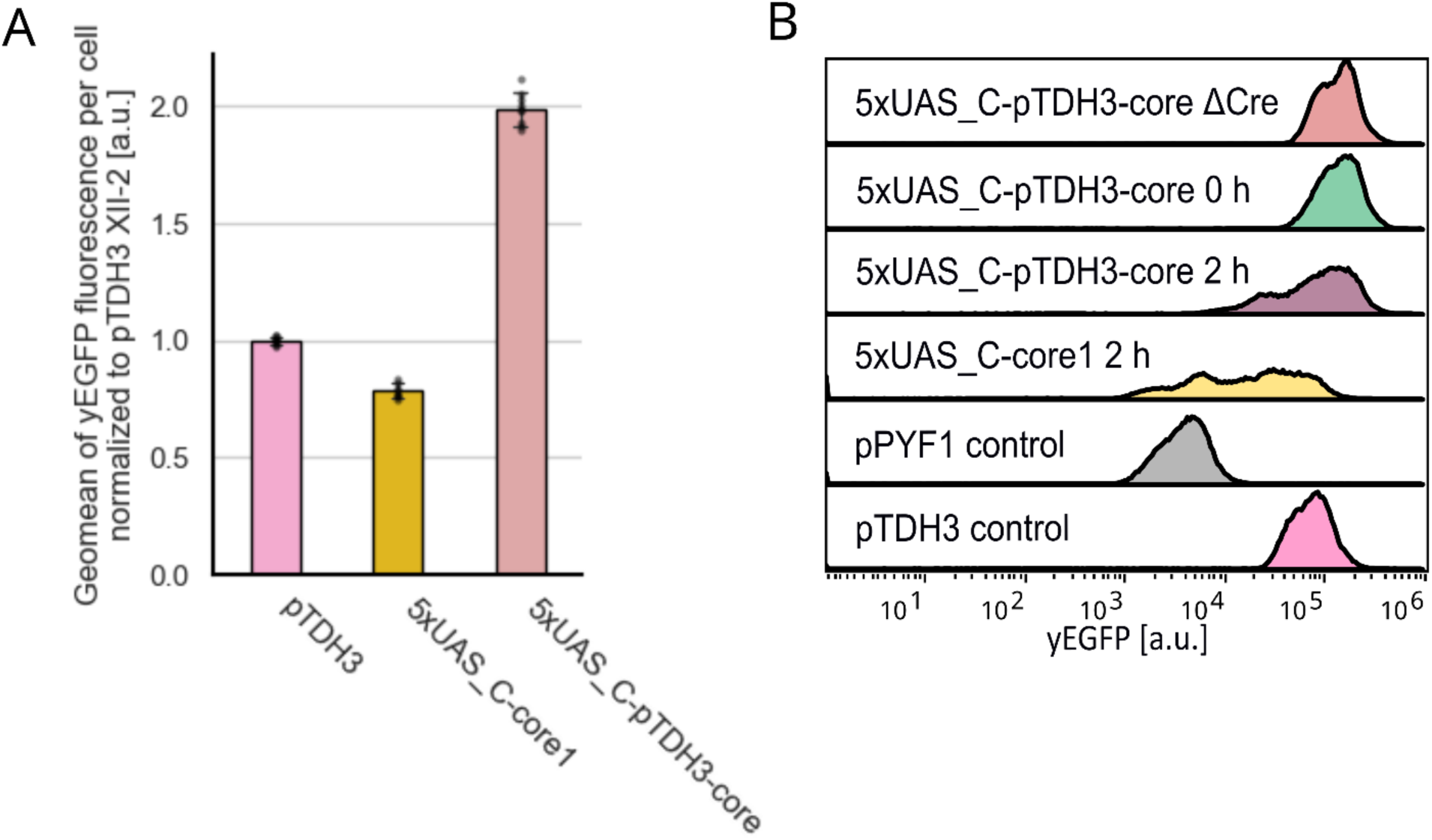
PULSE treatment of a 5xUAS-pTDH3-core promoter. (A) yEGFP fluorescence regulated by a 5xUAS_C-pTDH3-core promoter integrated into the XII-2 locus. The expression strength of this promoter even surpasses the *TDH3* promoter, showing twice its activity. yEGFP fluorescence was measured via flow cytometry in the exponential phase for three biological replicates with three technical replicates (indicated for each bar as gray dots). yEGFP fluorescence is normalized to the expression strength of the *TDH3* promoter. Error bars represent the standard deviation across all replicates. a.u., arbitrary units. (B) yEGFP fluorescence was measured for the Cre recombined (2 h red light) 5xUAS_C-pTDH3-core promoter cassette and compared to native promoter controls (pPYF1 and pTDH3), uninduced 5xUAS_C-pTDH3-core as well as the original Cre recombined 5xUAS_C-core1 promoter cassette. While the 5xUAS_C-pTDH3-core gives rise to a variety of strong promoters no weak promoters can be generated. yEGFP fluorescence was measured via flow cytometry in the exponential phase.

**Supplementary Figure 6.**
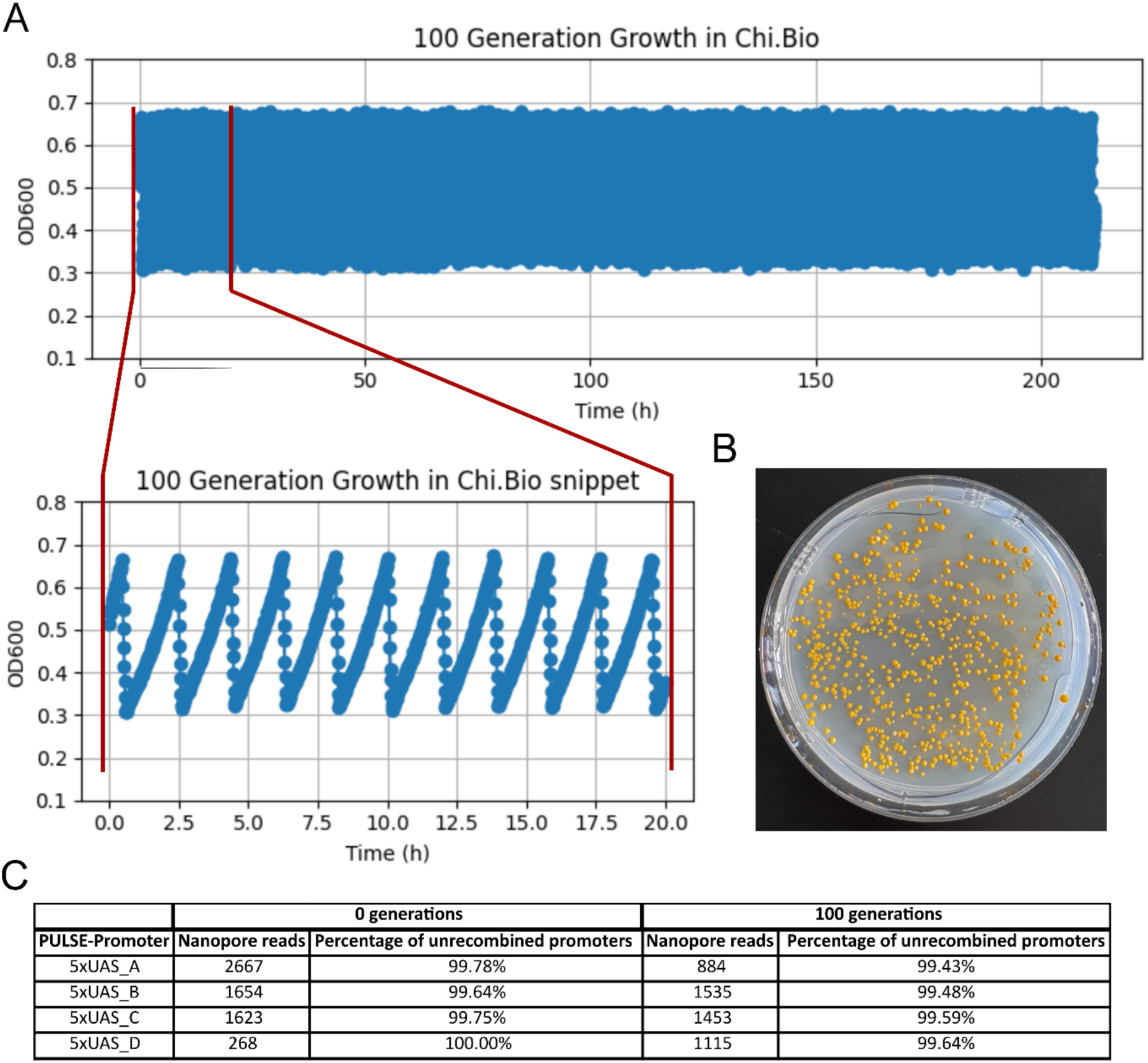
PULSE promoter stability in β-PULSE-0 over 100 generations of growth. (A) Growth curve of the β-PULSE-0 strain in the Chi.Bio turbidostat platform using the dither function. The culture was always diluted after one generation had passed over a time course of more than 200 h until 100 generations had passed. (B) Streaked out cells of the β-PULSE-0 strains after passing 100 generations. The similar color of the colonies hints on even expression levels of the pathway genes and therefore stable PULSE promoter cassettes. (C) Nanopore analysis of PCR amplified PULSE promoters before and after passing 100 generations. In both cases > 99% of the analyzed promoter sequences could be identified as the unrecombined 5xUAS PULSE promoters.

**Supplementary Figure 7.**
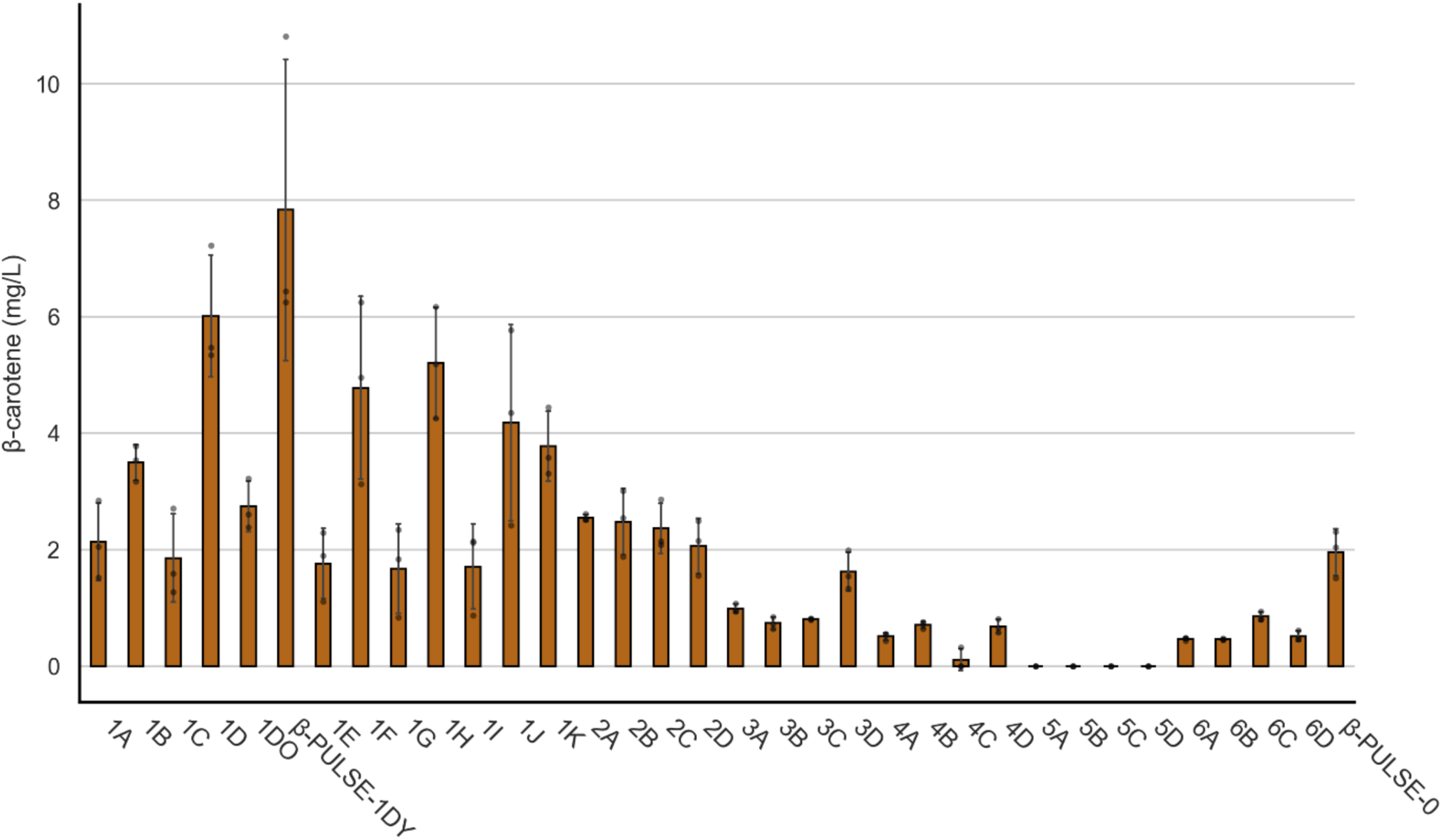
β-Carotene production of recombined β-PULSE strains. The recombined β-PULSE strains, as well as the β-PULSE-0 control where grown in 500 µl in a 48-well plate over a period of 72 h. β-Carotene was extracted and analyzed via HPLC. Strains with six different color phenotypes were compared to the control. The strains received a number based on their color (1- dark orange; 2- dark yellow, 3- yellow, 4- very faint yellow, 5- white and 6- slightly pink). The best producers were found analyzing the dark orange strains with β-PULSE-1DY as the best producer. The data was generated in one biological replicate with three technical replicates. Error bars represent the standard deviation across all replicates.

**Supplementary Figure 8.**
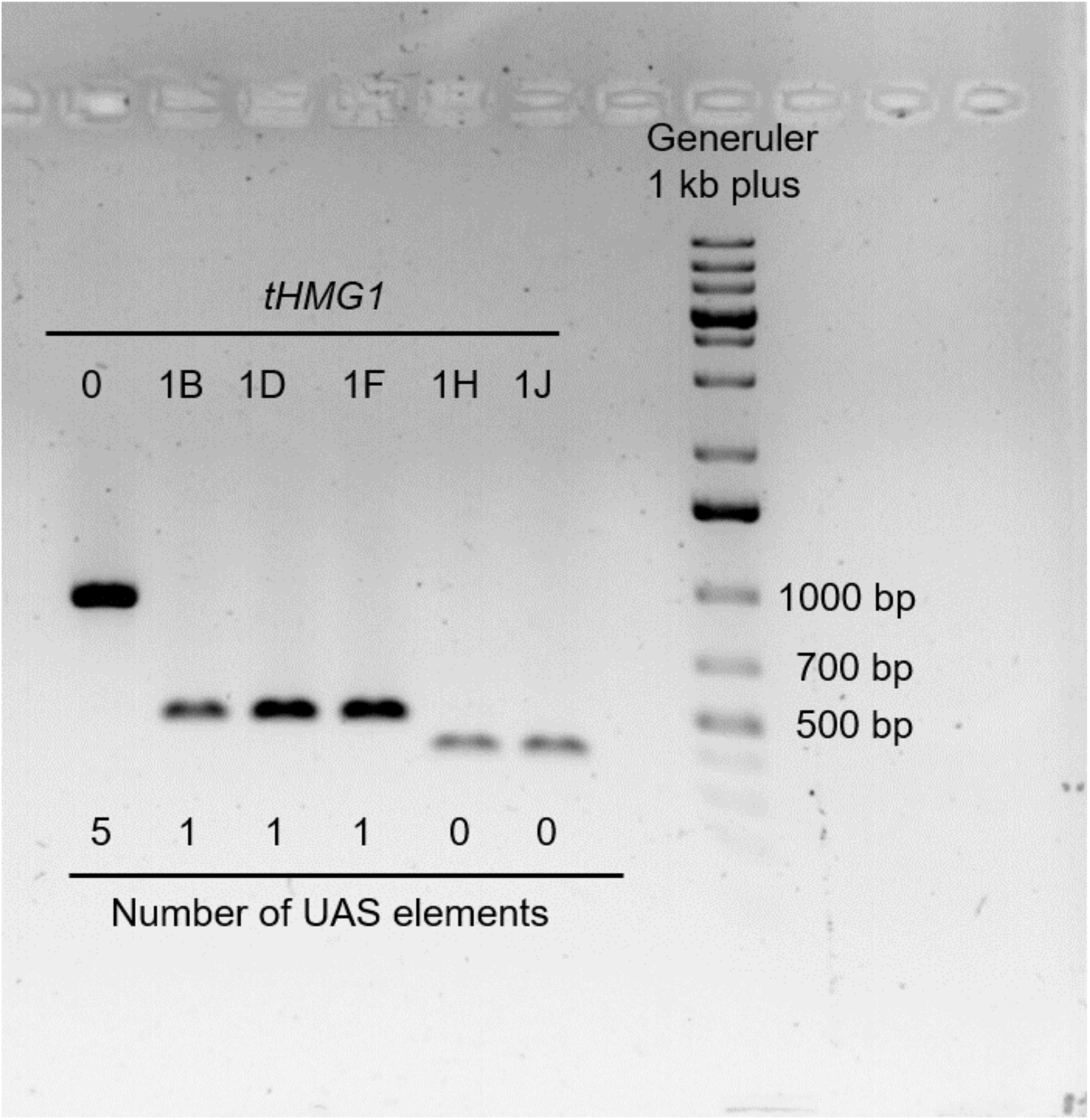
Analysis of *tHMG1* regulating promoters in the best β-carotene producers. Promoters of the β-carotene producers post recombination were amplified and analyzed on a 2% agarose gel compared to β-PULSE-0 (0). In all strains *tHMG1* is regulated by short promoters containing either one or no UAS element.

**Supplementary Figure 9.**
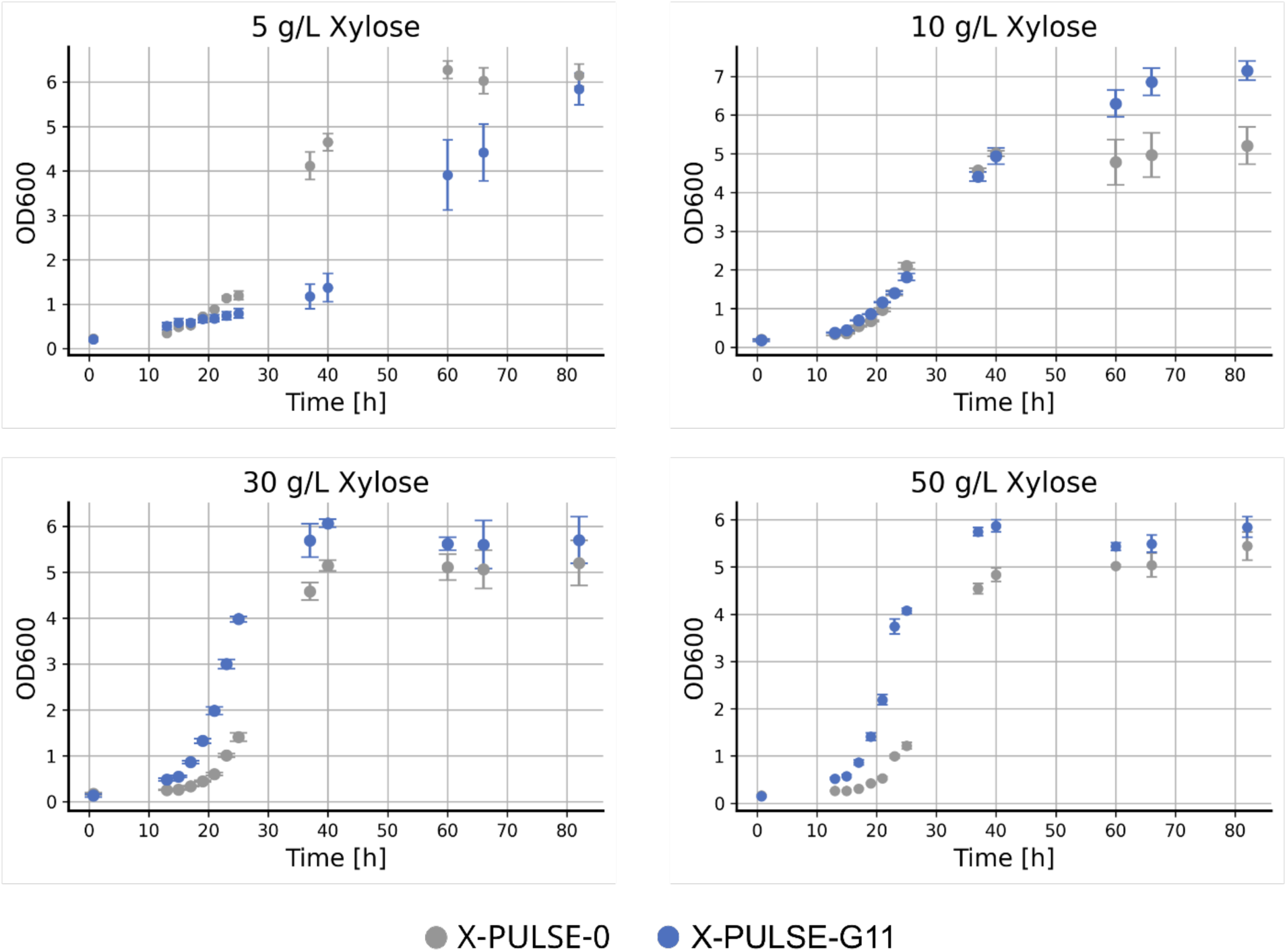
Growth in different xylose concentrations. Growth analysis of the optimized X-PULSE-G11 strain compared to the unrecombined X-PULSE-0 strain at different xylose concentrations in 48-well plates. X-PULSE-G11 grows slower at 5 g/L xylose concentration but outperforms X-PULSE-0 in the higher xylose concentrations. The data was generated for three biological replicates, each with three technical replicates. Error bars represent the standard deviation across all replicates.

